# Critical analysis of polycyclic tetramate macrolactam biosynthetic cluster phylogeny and functional diversity

**DOI:** 10.1101/2024.01.22.576670

**Authors:** Christopher P. Harper, Anna Day, Maya Tsingos, Edward Ding, Elizabeth Zeng, Spencer D. Stumpf, Yunci Qi, Adam Robinson, Jennifer Greif, Joshua A. V. Blodgett

**Affiliations:** Department of Biology, Washington University in St Louis. St Louis, MO; Columbia University School of Dental Medicine, New York, NY; Pfizer Inc, Chesterfield, Missouri; USDA-ARS, New Orleans, LA; Tulane School of Medicine, New Orleans, LA

**Keywords:** polycyclic tetramate macrolactams, phylogeny, biosynthetic gene clusters, comparative metabologenomics

## Abstract

Polycyclic tetramate macrolactams (PTMs) are bioactive natural products commonly associated with certain actinobacterial and proteobacterial lineages. These molecules have been the subject of numerous structure-activity investigations since the 1970s. New members continue to be pursued in wild and engineered bacterial strains, and advances in PTM biosynthesis suggests their outwardly simplistic biosynthetic gene clusters (BGCs) belie unexpected product complexity. Towards addressing the origins of this complexity and understanding its influence on PTM discovery, we engaged in a combination of bioinformatics to systematically classify PTM BGCs, and PTM-targeted metabolomics to compare the products of select BGC types. By comparing groups of producers and BGC mutants, we exposed knowledge gaps that complicate bioinformatics-driven product predictions. In sum, we provide new insights into the evolution of PTM BGCs while systematically accounting for the PTMs discovered thus far. The combined computational and metabologenomic findings presented here should prove useful for guiding future discovery.

**IMPORTANCE:** Polycyclic tetramate macrolactam (PTM) pathways are frequently found within the genomes of biotechnologically-important bacteria, including *Streptomyces* and *Lysobacter* spp. Their molecular products are typically bioactive, having substantial agricultural and therapeutic interest. Leveraging bacterial genomics for the discovery of new related molecules is thus desirable, but drawing accurate structural predictions from bioinformatics alone remains challenging. This difficulty stems from a combination of previously underappreciated biosynthetic complexity and remaining knowledge gaps, compounded by a stream of yet-uncharacterized PTM biosynthetic loci gleaned from recently sequenced bacterial genomes. We engaged in the following study to create a useful framework for cataloging historic PTM clusters, identifying new cluster variations, and tracing evolutionary paths for these molecules. Our data suggests new PTM chemistry remains discoverable in nature. However, our metabolomic and mutational analyses emphasize practical limitations to genomics-based discovery by exposing hidden complexity.

## INTRODUCTION

Polycyclic tetramate macrolactams (PTMs) are widely studied antibiotic-like bacterial natural products. They were first discovered in the bioactive fractions of microbial and marinelife extracts, but their biosyntheses remained uncharacterized until being linked to discreet biosynthetic gene clusters (BGCs) in *Lysobacter enzymogenes* C3 (1) and *Streptomyces* sp. strain SPB78 (2). Consequently, PTMs embody a relatively scarce example of an antibiotic family produced by both Gram-positive and Gram-negative organisms (3, 4). Over time, reports of PTM molecules and their encoding BGCs expanded beyond *Streptomyces* and *Lysobacter* to include several other filamentous actinomycetes (*Kitasatospora, Salinispora, Micromonospora, Saccharopolyspora, Saccharothrix, Umezawaea, Nocardiopsis, Actinokineospora,* and *Actinoalloteichu*s), gamma-proteobacteria (*Pseudoalteromonas, Colwellia, Gynuella, and Saccharophagus)*, as well as the firmicute *Pelosinus fermentans* (1, 2, 5–14).

Regardless of producer phylogeny, 66 all characterized PTM biosyntheses are initiated via an unusual hybrid-iterative polyketide synthase/ non-ribosomal peptide synthetase (iPKS-NRPS, encoded by *ftdB*, *ikaA,* and orthologs) that produces the polyene-tetramate precursor lysobacterene A (or related isomers) (15, 16). However, individual PTM biosynthetic pathways significantly diverge from this point, generally via combinations of oxidative decorations and carbocyclic ring arrangements (Fig 1A). There are now >85 known PTM congeners, and exploring PTM diversity has led to a growing body of valuable structure-activity relationship knowledge (17–19, 5, 20–22, 10, 23–26, 11). Testament to their structural diversity and their presumed interaction with a spectrum of biological targets, PTMs display a range of bioactivities. Examples include anti-inflammatory (27), antifungal (2), antibacterial (28), and predicted anticancer (29) activities, making the family compelling for future pharmaceutical and crop protective investigation. PTM BGCs are often targeted for synthetic refactoring to create new chemical matter or to probe their transcriptional regulation towards increasing production (10, 18, 22, 24, 30–32).

**Figure 1.**
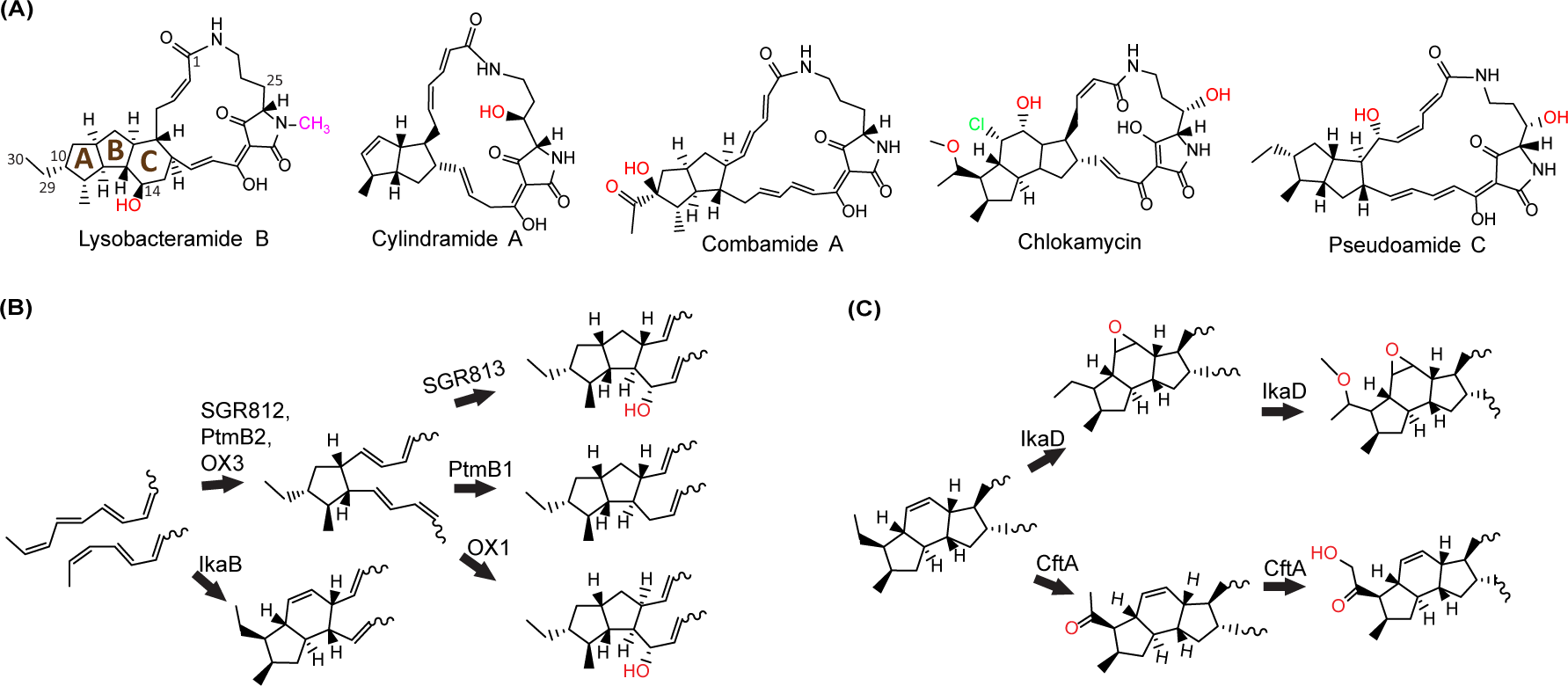
PTM structural diversity. (A) Example PTMs illustrating varying ring arrangements (brown numbers), tailored decorations (red, green & pink), and stereochemistry. (B) PTM PhyDHs catalyze an array of A and B ring arrangements. (C) PTMs sometimes involve multistep CYP450 oxidations, which are difficult to bioinformatically predict.

PTM BGCs typically feature relatively low gene content (3-7 genes), organized in operon-like arrangements clustered around their *ftdB* homologs. These features allow facile recognition in sequenced genomes via directed genome-mining or using targeted PCR screens in environmental isolates (6, 16, 33, 34). The ready availability of PTM cluster sequences has empowered numerous biosynthetic inquiries via classical genetic, biochemical, and synthetic-biological (6, 8, 14–16, 20, 22, 24, 34–38) approaches. As a result, several gross chemical features of PTMs (their polycycle patterns, in particular) are now largely predictable by BGC ORF content and gene identity (17). In brief, PTM cyclization cascades are driven by two classes of BGC-linked enzymes: flavin-dependent phytoene dehydrogenase homologs (PhyDHs) that form the A- and B-rings (e.g., IkaB, FtdC/FtdD, SGR813/SGR812, PtmB1/PtmB2, or OX1/OX2/OX3 (2, 8, 15, 32, 34)) and whenever present, nicotinamide-cofactor-dependent alcohol dehydrogenase (ADH) homologs (e.g., IkaC, FtdE and OX4) that form C-rings (39). Together, these two groups of enzymes are determinative for differentiating PTMs into polycycle type-classes. Illustrative PTM examples (Fig 1A) include combamide A with a 5/5 ring system (requiring two PhyDH homologs) (30), lysobacteramide B having a 5/5/6 ring system (requiring two PhyDHs and an ADH homolog) (15), and ikarugamycin-like chlokamycin which has a 5/6/5 ring system (requiring a single PhyDH plus an ADH homolog) (25). Finally, genes encoding cytochrome P450s (e.g., *ftdF, cftA, ikaD*, etc.) (2, 20, 26) and PTM C-25 oxidases (*ftdA* and homologs) (2, 14) are frequently associated with PTM BGCs where they direct oxidative decorations that diversify the class.

Despite the existence of >200 publications on PTM chemistry and biosynthesis, there remains sustained research interest in these molecules. This interest stems from several unusual features that have become hallmarks of the PTM family, including their atypical commonality and conservation across bacterial groups, the relative ease of their (bio)synthetic manipulation, their diverse bioactivities, plus their atypical enzymology. While core PTM cluster architectures are fairly conserved, an increasing number of PTM tailoring genes have been found outside the confines of their *ftdB*-centered BGCs (26, 40), which complicates product prediction. Finally, some PTM BGC-associated genes have biosynthetic functions that remain poorly predicted from sequence analysis alone (Fig 1B-C). Cytochrome P450s can perform not only regiodiverse oxidation chemistries, but some (e.g., IkaD & CftA) also catalyze multiple oxidation events, leading to unexpected shunt paths (Fig 1C) (6, 20, 26). Some PTM phytoene dehydrogenases are multifunctional too; certain B-ring forming PhyDH enzymes (e.g., FtdC, OX1/OX2) catalyze oxy-insertions while others (e.g., CmbB, PtmB1) apparently do not (Fig 1B) (35). Finally, several other PTM molecular features lack known enzymology, including the chlorine decorations of chlokamycin (18, 25), the terminal olefin of frontalamides (2), the apparent loss of backbone carbons in cylindramide and geodin (41, 42), and the oxidative decoration of PTMs whose BGCs lack CYP450s (18). Finally, because PTM BGCs continue to be encountered in newly sequenced genomes, frequent reassessment of the class becomes necessary.

Here, we describe the systematic classification of PTM BGCs from 302 bacterial strains to document the diversity and distribution of PTM BGCs. Our phylogenetic and genome mining efforts revealed new PTM cluster architectures are discoverable from local soil actinomycetes and indicate most PTM BGCs are vertically transmitted. Our gene-by-gene deconstructive approach was able to detect “hidden” diversity among seemingly syntenic BGCs, and in certain clusters it revealed surprising mosaicism that hints towards infrequent but important horizontal gene transfer. Finally, we demonstrated that predicting biosynthetic outcomes via bioinformatics alone remains challenging. This followed from BGC gene-reduction experiments in *Streptomyces* sp. strain JV180 and *Streptomyces olivaceus* NRRL B-3009, plus the comparison of several other wild strains. This work provides a useful map of systematically classified PTM clusters, while our PTM-targeted metabolomics exposes limitations in using BGC data to predict product diversity.

## RESULTS AND DISCUSSION

### Identifying PTM BGCs in previously-sequenced and newly-isolated environmental bacteria

Our searches for PTM BGCs focused on detecting FtdB orthologs because the enzyme is essential for all PTM biosyntheses, its polyketide synthase functions are different from all known type-I iterative PKSs, and FtdBs are readily differentiable from non-FtdB PKS-NRPSs *in silico* (6, 16, 33, 34). The relatively large size of FtdBs (3069-3354 aa; Table S1) lend themselves to well-resolved phylogenetic analysis (Fig 2, Fig S1). Our targeted searches to locate PTM BGCs in public repositories (see Methods) yielded 284 BGCs (after manual dereplication to prevent duplicate analysis, Table S2).

**Figure 2.**
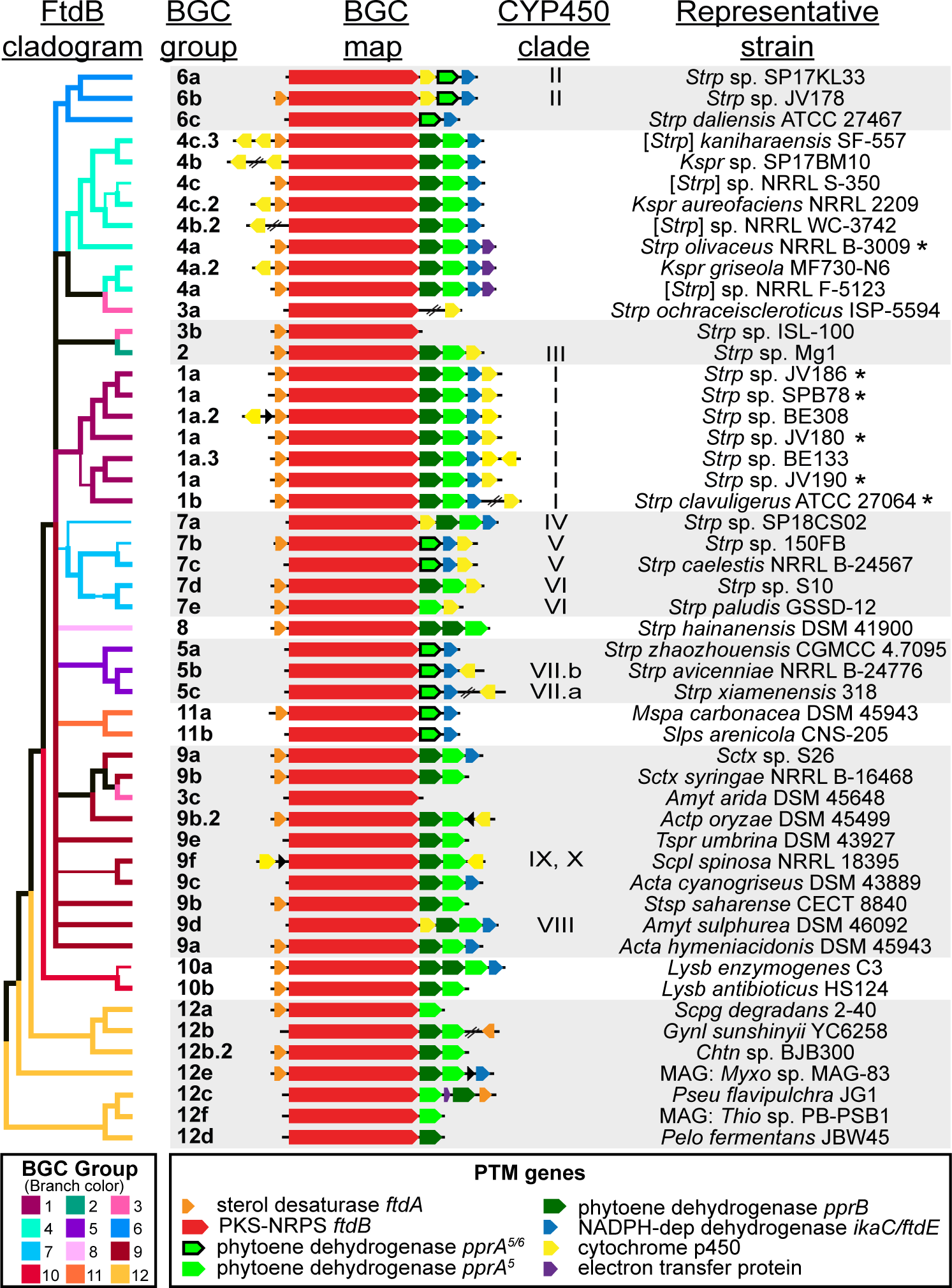
A FtdB- PTM BGC concordance. A cladogram representing 302 full-length FtdB sequences was collapsed for clarity and linked to a meta-analysis of associated PTM BGC types, representative strains, and CYP450 clades (see Figs S1 and Fig S3 for associated maximum-likelihood phylogenies). Note representative strains for each architecture are shown; this is not indicative of complete host taxonomy for each BGC type (See Fig S2 for detail and Table S2 for abbreviations). Genus names with brackets indicate a potential taxonomic discrepancy with the current naming convention. Asterisks indicate strains examined via PTM-targeted metabolomics in this study.

We surmised some PTM BGC types would be rare or under-represented in public databases. To create a more diverse study set, we leveraged an *ftdB*-targeted degenerate-PCR screen (6) to identify PTM BGCs in actinobacterial isolates sourced from local soils. Based on amplicon sequence divergence or perceived scarcity, we selected 18 strains for complete cluster analysis via whole-genome sequencing (see nucleotide accessions). Along with complete PTM BGCs, this yielded *atpD, recA, rpoB,* and *trpB* sequences for host multi-locus (MLSA) phylogeny. Thus, public and environmental sequences yielded 302 complete PTM BGCs for comparison.

In terms of producer phylogeny, these efforts immediately expanded the catalog of putative PTM producers into the actinobacterial genera *Actinophytocola*, *Amycolatopsis, Thermomonospora,* and *Streptosporangium.* Several new proteobacterial producers were also identified including *Chitinimonas*, plus several metagenome assembled genomes (MAG) including *Cellvibrio* from citrus rhizosphere (43)*, Thiohalocapsa* from a ‘pink berry’ sulfur cycling consortium (44–46), and a putative Myxococcales bacterium found in wastewater sludge (47) (Fig S2).

### A deconstructive approach to PTM BGC cataloging

Since the first PTM BGCs were sequenced >10 years ago, efforts to categorize their BGCs have been largely descriptive, guided mostly by synteny and the presence/absence of key biosynthetic genes. In pursuing a systematics-driven analysis of PTM BGC diversity, we assumed synteny-based binning of BGCs would eventually become limiting. This is because certain PTM-linked enzymes (especially cytochrome P450s and phytoene dehydrogenases) catalyze diverse reactions and are thus key drivers of PTM structural diversification (especially in combination). However, these enzymes are difficult to assign accurate functions to, especially in the absence of individual enzyme phylogenies. Accordingly, organizing PTM BGCs into discrete cluster types required particular attention to associated CYP450s (6, 20, 22, 26) and PhyDHs (8, 15, 18, 24, 34, 35, 48). We also avoided the use of concatenated trees (sometimes used to characterize other antibiotic BGCs (49, 50)) due to the high risk of conflating similarly organized but functionally different PTM loci. Thus, maximum-likelihood (ML) phylogenies of individual PTM-encoded enzymes (Fig 3, Fig S1, S3-5) were used for accurate, clade-based systematic binning of PTM enzymes and the resulting enzyme histories were used in-composite to identify meaningful enzyme sequence differences and evidence for BGC mosaicism.

**Figure 3.**
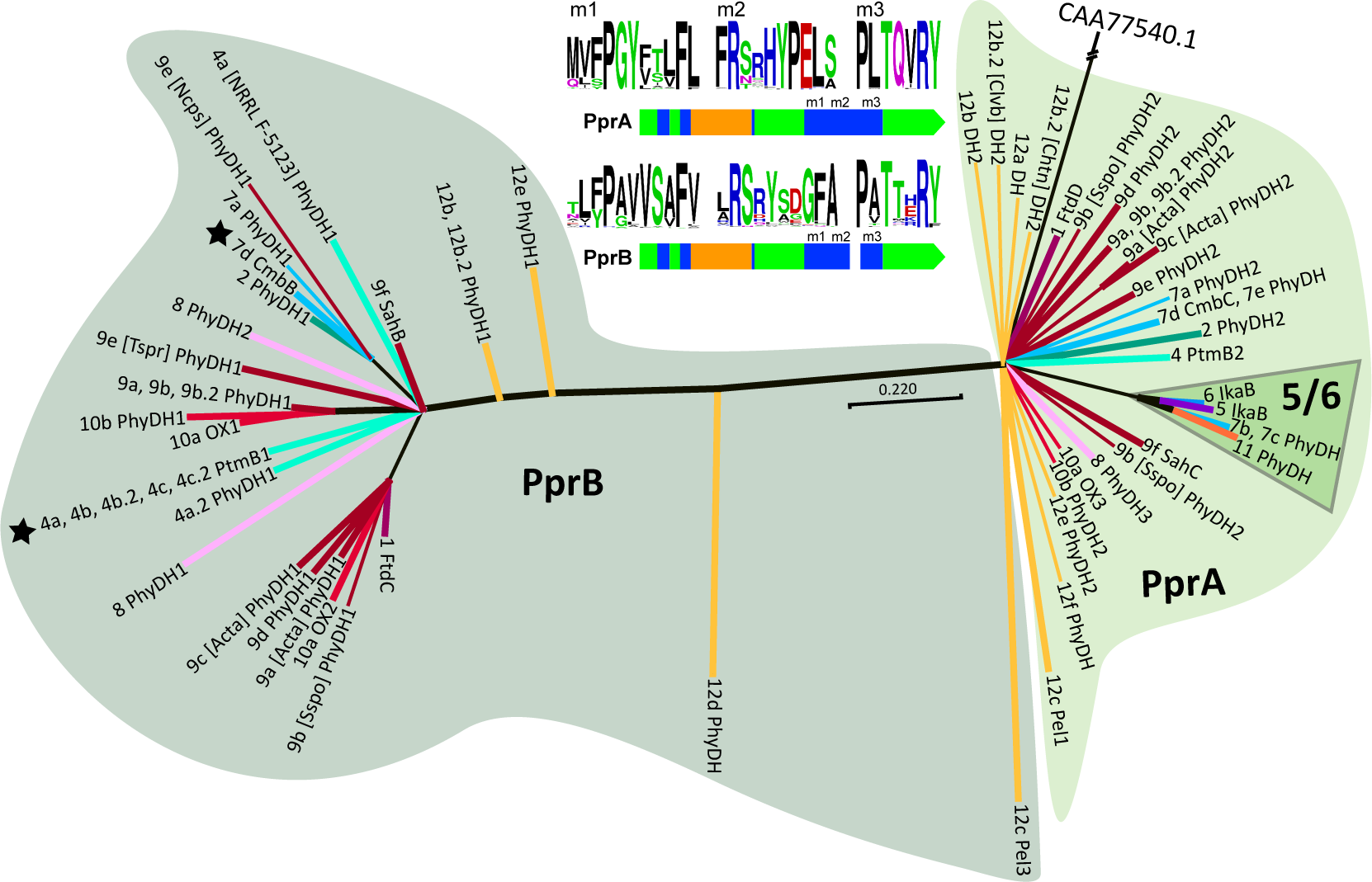
PTM- associated PhyDH ML phylogeny and sequence analysis assist polycycle prediction. Branches are colored according to BGC groups, UF-Bootstrap threshold is 95%, and branches >99% are bolded. Stars denote experimentally characterized non-oxygenating PprBs. Protein domains are colored according to *Pantoea ananatis* CrtI homology (51), where green is FAD-binding, blue is substrate-binding, and orange is a non-conserved membrane helical region. Web logos illustrate amino-acid motifs differentially enriched in PprA or PprB-type PhyDHs.

### PTM tailoring enzyme phylogenies

PTM-associated phytoene dehydrogenase-like (PhyDH) enzymes are named for homologs in carotenoid biosynthesis (CrtI (51)), but despite their specialized roles in PTM polycycle formation (34) (Fig 1B), the literature has lacked a standard, function-based nomenclature for these enzymes. Instead of continuing historical naming conventions (e.g., OX1, OX2 and OX3; PhyDH1, PhyDH2, etc.), we adopted a phylogeny and function-based nomenclature rooted in the distinct catalyses of **P**TM **P**hyDH-like **R**ing-forming enzymes (Ppr’s). Accordingly, the two PhyDHs required for the stepwise formation of a 5/5 ring PTM from FtdB products were termed PprA^5^s (initial cyclopentenyl, A-ring formation) and PprBs (for B-ring formation). In contrast, forming 5/6 polycycles (34) requires a single specialized PhyDH we termed PprA^5/6^. Our Ppr ML phylogeny (Fig 3) multifurcates, with PprA^5^s forming discrete clades congruent with FtdB phylogeny. However, a distinct branch emerging from the Ppr polytomy reveals that all known PprA^5/6^ proteins (plus several synteny-suggested ones) are monophyletic. Interestingly, this suggests PprA^5/6^s arose from ancestral PprA^5^s, suggesting 5/6/5 PTMs may have evolved after their 5/5/6 counterparts.

In contrast, PprBs branch deeply to form several subclades. Finding well-delineated PprB subclades initially suggested the possibility of ascribing precise catalytic function (oxygenation status, in particular) via conservation with literature-characterized members. However, we found non-hydroxylating and hydroxylating PprBs to be polyphyletic (Fig 3), preventing us from building on this idea and contrasting with earlier work in this vein (35). Our PhyDH analyses revealed other insights that may eventually inform catalysis. Most PprAs were distinguished from PprBs by a ∼19 aa indel and CrtI homology suggests this region lies within a substrate-binding domain (51). This suggests a basis for differential substrate recognition and serves as useful landmark for parsing PprAs and PprBs from one another. However, we found intriguing exceptions to this pattern in PhyDHs extracted from PTM group 12c (Alteromonadales). Here, the associated PprAs and PprBs both contain a distinct sequence insertion from all other Pprs (likely contributing to their PhyDH-clade resolution) (Fig 3). Interestingly, the 12c BGC *ppr* genes are arranged in reverse to all other known PTM BGCs (Fig 2). Recent analyses of Pel3, a 12c PprB, showed catalysis of a rare C6 hydroxylating, C7-C14 crosslinking B-ring variant during heterologous pseudoamide biosynthesis (48). These observations suggest an atypical evolutionary history for some Pprs, and the need for a more rigorous way of differentiating PTM PhyDHs beyond the indel region. Leveraging our PprA and PprB deep sequence comparisons, we revealed differentially enriched sequence motifs that should further assist classification efforts (Fig 3, Weblogo insets).

PTM-associated CYP450s have been intensely studied for understanding PTM oxidative decorations and exploiting the enzymes for molecular diversification. Our CYP450 phylogeny was created from an alignment of 232 CYP450s extracted from within the operon-like arrangements of core PTM BGCs, and those encoded nearby (within 10 kb) these clusters. The decision to include the latter group arises from increasing numbers of PTM-active enzymes (i.e., CYP450 IkaD (26) and HSAF *N*-methylase (40)) that locate outside of core BGCs, but are sometimes found nearby. Linking these unclustered CYP450s to PTM catalysis will require future experimentation so clade-numbers were not assigned to these particular CYP nodes (Fig S3). The resulting CYP450 ML phylogeny has two main branches, one comprised of CYP107-family enzymes (FtdFs) linked to Group 1 PTM BGCs (**I,** SGR and frontalamide-type PTMs) (2, 22, 23) plus multiple uncharacterized CYP450s upstream of Group 4 BGCs in certain *Kitasatospora*. The other main branch featured several subclades of CYP107 P450s with known biosynthetic roles, including CftA (**II**, clifednamides) (6, 20), IkaD (**VII.a**, ikarugamycin epoxide) (26), Mg1D (**III**, combamides B and G) (22), and CbmD (**VI**, combamides) (30). Other prominent subclades featured characterized CYP105 family members (**IX**, **X**; SahD and SahE of sahamide biosynthesis) (10), yet-uncharacterized CYP450s likely acting in PTM biosynthesis due to gene clustering (**IV**, **V**, **VII.b**, **VIII**), plus BGC proximity-flagged ones (e.g., BE133 CYP(2)). Finally, some *Kitasatospora* CYP450s [*Kspr* CYP(2)] upstream of Group 4 BGCs were considerably diverged from other members of this branch.

FtdAs (PTM C-25 hydroxylases, Fig S4) and FtdEs (alcohol dehydrogenase-like C-ring forming enzymes, Fig S5) are critical for PTM structure determination and bioactivity (2, 39). In contrast to PhyDHs and cytochrome P450s, the catalyses performed by FtdAs and FtdEs are understood to be fairly conserved (14, 18). This suggests against them having pathway-specialized roles. The phylogenies of both enzymes appeared largely concordant with FtdB (Fig S1), suggesting sequence diversity seen in these proteins likely reflects drift and host speciation rather than catalytic diversity. This idea was supported by MLSA phylogeny of concatenated host housekeeping genes (Fig S2), where the most divergent FtdBs (an enzyme of highly conserved function) correlated with the most distant genera. Therefore, when interpreting PTM BGCs, we surmised *ftdA* and *ftdE* presence or absence should be emphasized over the low possibility of catalytic divergence.

### A combination of FtdB phylogeny and BGC synteny is useful for estimating PTM diversity

Microbial genome-mining requires the accurate recognition of BGCs and identification of novel clusters. To do this for PTMs, a combination of evidence including traditional synteny, individual enzyme phylogenies and published PTM genetic, biochemical, and structural data were used where possible. BGC types identified using these data were anchored to an FtdB cladogram, resulting in a map that reflects both sequence relatedness and differential enzyme content (Fig 2). Alpha-numeric identifiers were used to describe groups of BGCs according to their FtdB clades, and BGC variations within those clades. The first digit indicates FtdB branch (with the exceptions of Groups 3, 9 and 12, where groups of divergent-but-similar BGCs were binned). The letter code indicates plasticity among the core PTM genes, and decimals indicate potentially important tailoring genes are encoded nearby (generally CYP450s).

PTM are divisible into two major families based on their polycycle patterns and the specialized enzymology associated with each. One is the 5/6/5 PTMs and the other is comprised of the 5, 5/5, and 5/5/6 PTMs. Assigning BGCs to the first one was straightforward using the diagnostic presence of a PprA^5/6^ (but no PprB), plus an ADH. BGCs having these hallmarks were found in PTM groups 5, 6, and 11 (Fig 2) and distributed among diverse actinobacterial hosts. Group 5 clusters (5 a-c) were found in a distinct clade of *Streptomyces* spp (Fig S2) and include known loci for ikarugamycin and its oxidized analogs (capsimycins via IkaD, a type **VII.a** CYP450) (26). Interestingly, the *ikaD*s of Group 5 strains all locate downstream of their core ikarugamycin-type BGCs, but show highly variable distances between them (also observed in (37)). Group 6 BGCs were mostly found in *Kitasatospora* and *Streptomyces* thought to be plant-associated (Fig S2) (52–58). Their BGCs include several clifednamide loci (type **II** CYP450s, CftA) that include (6b) or lack (6a) *ftdA*s (6, 20). Interestingly, the *S. daliensis* group 6c BGC lacks a clustered *cftA*, thus likely producing ikarugamycin instead of clifednamides. Finally, the Group 11 BGCs from *Salinispora arenicola* and *Micromonospora* spp. (Fig S2) lack CYP450s. Some members produce ikarugamycin (8) or butremycin (9), presumably via cluster types 11b and 11a respectively.

In contrast with 5/6/5 PTMs, predicting the functions of the remaining PTM clusters is more complex. 5/5/6 PTM carbocycles are produced stepwise, with ring A5 first (PprA^5^), then B5 (PprB), then C6 (ADH). Thus, based on several biosynthetic studies (reviewed (17)), BGCs lacking any one of these enzymes must direct lower (or no) polycycle complexity. Thus, the simplest BGC architectures we encountered should produce otherwise-acyclic tetramate polyenes similar to lysobacterene A (i.e., Group 3a (CYP450 adjacent), b (*ftdA*-adjacent), c (*ftdB*-alone)), molecules that are technically not polycyclic macrolactams. Such compounds have never been found as native pathway products. As with most BGCs analyzed here, it remains to be determined if these loci are active or are past relics of genome rearrangement.

PTMs having a single five-membered (A5) carbocycle also remain undiscovered from nature, but have been produced in engineered systems (e.g., lysobacterene B (15), pactamide E (18, 24), sahamides A-D (10)). We identified potential native producers of such compounds among the non-actinobacterial group 12 BGCs (Fig S2, Table S2). *Saccharophagus degradans* (12a) and *Thiohalocapsa* sp. PB-PSB1 (12f) could produce PprA-dependent A5 ring PTMs, while *Pelosinus fermentans* (12d) seems to encode an unprecedented B5 PTM via its lone PprB.

5/5-polycyclic PTMs are produced by natural and pathway-engineered bacteria (10, 12, 13, 15, 30, 48, 59, 60). In native strains they can arise from dedicated pathways, but they are also associated with incomplete 5/5/6 PTMs biosynthesis (31, 32, 61, 62). When mining for dedicated pathways, multiple clusters lacking adjacent ADH homologs and encoding conserved PprA^5^ and PprBs were detected. *Streptomyces* group 2 BGCs have a clustered *ftdA* and clade **III** CYP450, while the lone group 8 BGC encodes one PprA and two PprB homologs. Group 8 *pprB* multiplicity is unusual and was here-to-fore undocumented in actinomycetes, but it is similar to that of Gram-negative *Lysobacter enzymogenes* C3 (Group 10a). In *L. enzymogenes*, PprB multiplicity drives 5/5/6 polycycle stereo-diversification (15), which could be tested in Group 8. Interestingly, *Lysobacter antibioticus* (Group 10b) also appears to encode a dedicated 5/5 PTM pathway; it lacks one of the PprB homologs and the ADH of *L. enzymogenes*. Multiple other Gram-negative strains encode similar BGCs, including groups 12b and 12c. Finally, Group 9 PTM BGCs of the Nocardiaceae and Pseudonocardiaceae (Fig S2, Table S2) encode several inferred 5/5 PTM pathways (*ftdA*-associated 9b, 9b.2; *ftdA*-minus 9e, 9f).

5/5/6 PTMs were well-documented from a diversity of bacterial producers in the pre-genomic era (61, 63, 64). Several Group 1 BGCs have since been linked with 5/5/6 PTM production. Group 1 BGCs are exceptionally well-represented in genome databases, comprising >50% of our dataset (Fig S1). Perhaps unsurprisingly, the Group 1 BGC architecture was among the first identified during early genome-mining efforts, and 5/5/6 PTMs (and certain biosynthetic intermediates) have been isolated from *Streptomyces* (Fig S2) scattered across multiple Group 1 FtdB subclades. Known Group 1 PTM producers include *Streptomyces* sp. JV180 (maltophilin-like congeners) (38), *S. griseus* IFO13350 (maltophilin-like, and alteramide-like congeners) (24, 32, 38), *Streptomyces* sp. SCSIO40010 (10-epi-maltophilin) (23), multiple strains of *S. albidoflavus* (somamycins, dihydromaltophilin [HSAF], 10-epi-HSAF, xanthobaccin C, and alteramide stereoisomers) (19, 31, 65, 66), and *Streptomyces* sp. SPB78 (frontalamides) (2).

However, 5/5/6 PTM production is not limited to Group 1. Some bacteria having groups 9a, 9c, and 10a BGCs produce 5/5/6 PTMs identical to certain Group 1 products (7, 24, 64, 67). Interestingly, Group 9a, 9c, and 10a BGCs are similar in terms of gene content but are distributed among highly divergent genera (*Streptomyces*, *Actinoalloteichus*, *Saccharothrix*, *Actinokineospora*, and *Lysobacter*). Most of their known products have a hallmark hydroxyl or carbonyl group on C14. Previous works suggest hydroxylation at this position is PprB-catalyzed in Groups 1 and 10 (i.e., FtdC, OX1, OX2), so we surmise analogous catalyses occur in Groups 9a and c (whose PprBs clade with *Lysobacter* OX1 and OX2). However, the mechanism underlying C14 oxidation to a carbonyl remains unknown.

*Streptomyces* and *Kitasatospora* Group 4 BGCs appeared similar to Group 1 5/5/6 clusters but lacked clade **I** CYP450s. Further, several members (Group 4a, a.2, c) encoded a short gene (*ftdG*) for a Rieske 2Fe-2S-cluster protein, but this was not definitive for the group (4b.2, 4c.2, 4c.3). Group 4 PTM production has never been examined in native strains, but reconstitution of 4a BGC (excluding *ftdG*) from *Streptomyces* sp. SCSIO 02999 in *Streptomyces lividans* TK64 resulted in the discovery of 5/5/6 pactamides (18) and revealed the existence of C12-non-hydroxylating PprBs. This critically differentiates SCSIO 02999 core PTM biosynthesis from other 5/5/6 PTMs (18). Because Group 4 BGCs lack obviously clustered CYP450s but some pactamides feature oxidative decorations on their polycycles (18), the need for unclustered oxidases is implicated. This highlights CYP450s in the genome neighborhoods of BGCs 4a.2, 4b, 4b.2, 4c.2, 4c.3 for possible involvement (Fig S6). Finally, the biosynthetic role of *ftdG* in Group 4 BGCs remains enigmatic. Initial studies disregarded it as belonging to adjacent xiamycin BGCs (18, 68), but in SCSIO 02999 we noted the start codon of *ftdG* likely overlaps with the stop codon of upstream *adh*, which is common in operons. Furthermore, Group 4a.2 *Kitasatospora* BGCs feature *ftdG*s not juxtaposed with any xiamycin locus, again suggesting *ftdG*s are PTM-associated.

### Group 7 PTM BGCs show unusual BGC diversification and mosaicism

Group 7 stood out against all other PTM BGC groups in our analysis because it includes five different BGC architectures linked to four different carbocyclic ring patterns (5, 5/5, 5/6/5 and 5/5/6; Fig 4). Group 7 BGCs are carried by several taxonomically disparate strains of *Streptomyces* (Fig S2), and our use of individual protein phylogenies revealed evidence of unusual mosaicism in the group. For example, group 7 FtdAs (where present) and FtdBs clade similarly between their respective ML phylogenies, but the PprA^5/6^ genes of Groups 7b and 7c appear most closely related to those of group 11 from Micromonosporaceae (Fig 3), suggesting gene flow from outside of the *Streptomyces* genus contributed group 7 BGCs for 5/6/5 PTMs. In contrast, Groups 7a and 7d harbor *pprA^5^* and *pprB*s, predicting 5/5 and 5/5/6 PTM products, while the sole Group 7e BGC from *S. paludis* contains only a *pprA^5^*, thus putatively encoding PTMs with an A5 ring PTM. Interestingly, group 7a (5/5/6 carbocycles) and 7b-c (5/6/5 carbocycles) FtdEs are most related to those of other 5/6/5 clusters (IkaC-branch, Fig S5), again highlighting atypical gene patchworks in these BGCs.

**Figure 4.**
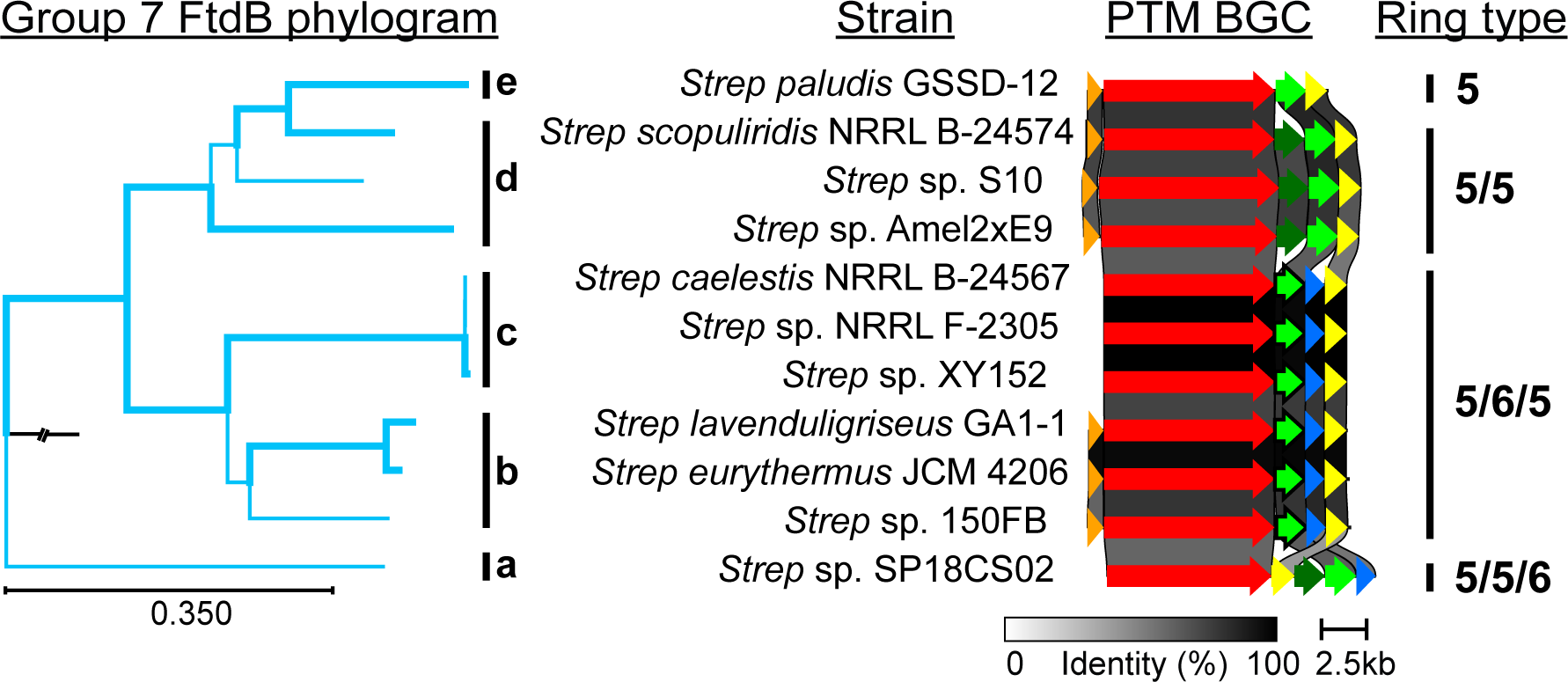
Group 7 BGCs are predicted to encode for an atypical diversity of PTM structures. FtdB protein phylogeny indicates the core biosynthetic enzymes of Group 7 BGCs are closely related, which starkly contrasts with the diverse tailoring gene content seen among individual BGC types. This suggests an unusual and exceptionally diverse array of PTM scaffolds are encoded by Group 7, compared to all other BGC groups identified in this work.

There was also remarkable plasticity in the CYP450 gene positions and their assigned clades in group 7. The CYP450 (**IV** clade) of group 7a CYP450 is encoded directly downstream of *ftdB*, a gene arrangement usually associated with 5/6/5 clifednamide BGCs (Group 6a/b). In turn, this unusual cluster outwardly appeared identical to the Group 9d BGC of *Amycolatopsis sulphurea* (Fig 2), but these clusters are non-orthologous because the latter has a clade **VIII** CYP450 (Fig S3). Group 7a and 9b cluster architectures are both new to the PTM literature, with the former being discovered in a streptomycete isolated from our local soils. In contrast with 7a, the remaining group members have CYP450s (clades **V** and **VI**) encoded downstream of their core biosynthetic genes. Most of the BGCs in group 7 remain experimentally uncharacterized, but the *Streptomyces* sp. S10 group 7d cluster was interrogated via synthetic biology to produce combamides (5/5 PTMs oxidatively tailored via clade **VI** CYP450 CmbD, Fig 1A, and Fig S3 (30)).

### PTM biosynthesis appears to have a complex history

Vertical inheritance is argued to be the main mechanism of natural product BGC evolution, because on evolutionary timescales horizontal gene transfer (HGT) of large clusters is likely infrequent (69). However, it is also recognized that BGCs and the genes that comprise them can have individual evolutionary paths (even within BGC families (70)). Some PTM BGC types showed clear patterns of vertical transmission. For example, Group 5 clusters were found only within a particular monophyletic clade of *Streptomyces* spp (Fig S2). However, for many other PTM groups, such clear host -BGC concordance was absent. Furthermore, our Group 7 analysis provided compelling evidence for HGT, albeit for a subset of genes (Fig 4) rather than a complete BGC. Among the Group 7 BGCs is a putative ikarugamycin cluster that, remarkably, may have arisen from gene exchange into an ancestral 5/5/6 locus (based on nearest neighbors). It seems ikarugamycin BGCs may be particularly subject to HGT, owing to their broad stochastic distribution among actinomycete hosts and their linkage with multiple FtdB clades (Fig 2). Finally, we noticed that at least ten PTM BGCs are adjacent to transposase genes, including members of Group 1, 4, 6, 9, and 12. It is intriguing as to what role these may have had in the spread of PTM BGCs. In sum, our analysis of FtdB and host phylogenies suggest an evolutionary history involving both vertical transmission and HGT.

### New FtdB clade-specific probes are necessary capture PTM diversity

While less common since the advent of inexpensive genome sequencing, degenerate PCR remains a powerful approach for screening environmental microbes for genes and pathways of interest. We have isolated several PTM producing actinobacteria this way, using primers YQ158/YQ159 that target *ftdB*s ((6) and this study). However, those primers were designed when far fewer FtdB sequences were available and based on the much larger FtdB sequence library presented here, we realized new tools would be required to access the full diversity FtdB sequences. Accordingly, we used a phylogeny-guided degenerate oligonucleotide design strategy (see Methods, Table S3) and the resulting primers were validated against a panel of PTM^+^ bacteria having diverse FtdB sequences (Fig S7). This resulted in new primer sets that facilitated more robust cross-clade *ftdB* amplification than YQ158/159 (i.e., MTftdB1B, MTftdB4, MTftdB3B), while others gave robust amplification of certain *ftdB* sequences that were difficult to amplify with other primers sets (such as MTftdB6 for *Streptomyces* sp. SP18CS02 and MTftdB7 for *Pseudoalteromonas luteoviolacea*). Together, these new tools embody a robust toolset for the targeted discovery of PTM BGCs from environmental bacteria. Finally, sequence fragments of BGCs can be used to extrapolate potential BGC novelty, which greatly empowers genome mining (i.e., KCDA genes in (71)). To test if the sequences of our new *ftdB* amplicons might predict PTM BGC novelty, a bank of degenerate-primer delimited amplicons was created in silico, and their in-frame translation products were used to build a ML phylogeny (Fig S8). This “amplicon tree” largely recapitulated the topology of the full-length FtdB tree (Fig S1, Fig S8), with some expected loss of resolution. Thus, FtdB amplicon phylogeny can be used to estimate PTM BGC novelty, within the constraints revealed by our FtdB-BGC concordance (Fig 2).

### Using Group 1 PTM-targeted metabolomics to assess the fidelity of gene-to-molecules prediction

We previously showed actinobacterial strains harboring identical PTM clifednamide BGCs can display a range of congener titers, distributions, and types (6). Group 1 PTM BGCs had yet to be systematically surveyed in this way, and we thus asked how well PTM locus conservation translates into conserved product profiles, especially given how much FtdB diversity is represented within Group 1. To sample PTM Group 1 product diversity across species, we leveraged established PTM-targeted metabolomics methods to compare five strains that span the group’s the FtdB diversity: *Streptomyces* sp. JV180, *Streptomyces* sp. SPB78, *Streptomyces* sp. JV186, *Streptomyces* sp. JV190, and *Streptomyces clavuligerus* ATCC 27064. Organic extracts from spent culture media of each were analyzed via high pressure liquid chromatography-tandem mass spectrometry (HPLC-MS/MS), with precursor ion scans for *m/z* 139.2 or 154.2 differentiating 25-deOH and 25-OH PTM congeners (38). The *m/z* transitions for each putative PTM are listed in Table S4, and product ion scans were performed for the most abundant base masses. The resulting MS^2^ spectra were subsequently used to differentiate mass-similar features via fragment signatures, and to compare these against known PTMs to assign membership to the PTM family. All putative PTMs were orthogonally identified by comparing product profiles against published Group 1 null mutants of *Streptomyces* spp strains SPB78 and JV180.

We found all tested group 1 strains produced highly similar PTM profiles (Fig 5A, Fig S9A. Specifically, the tested strains all produced a PTM (**1**) that is possibly identical to biosynthetic intermediate FI-2 of SPB78 based on retention time, mass transition, and fragmentation pattern (Fig 5B), plus another PTM (**13**) had high similarity to a maltophilin authentic standard (Fig S9A-B). In addition, several other commonly distributed putative PTM signals (**2**, **5**, **6**, **8**, **10**, **12-16**) were revealed by comparison against from PTM null mutants. Furthermore, two minor peaks were shared by only a subset of strains: **3** in JV180, JV186, and JV190, and **11**, found in SPB78 and ATCC 27064. Finally, SPB78 generated multiple minor PTM signals not seen in other strains (**7**, **9**).

**Figure 5.**
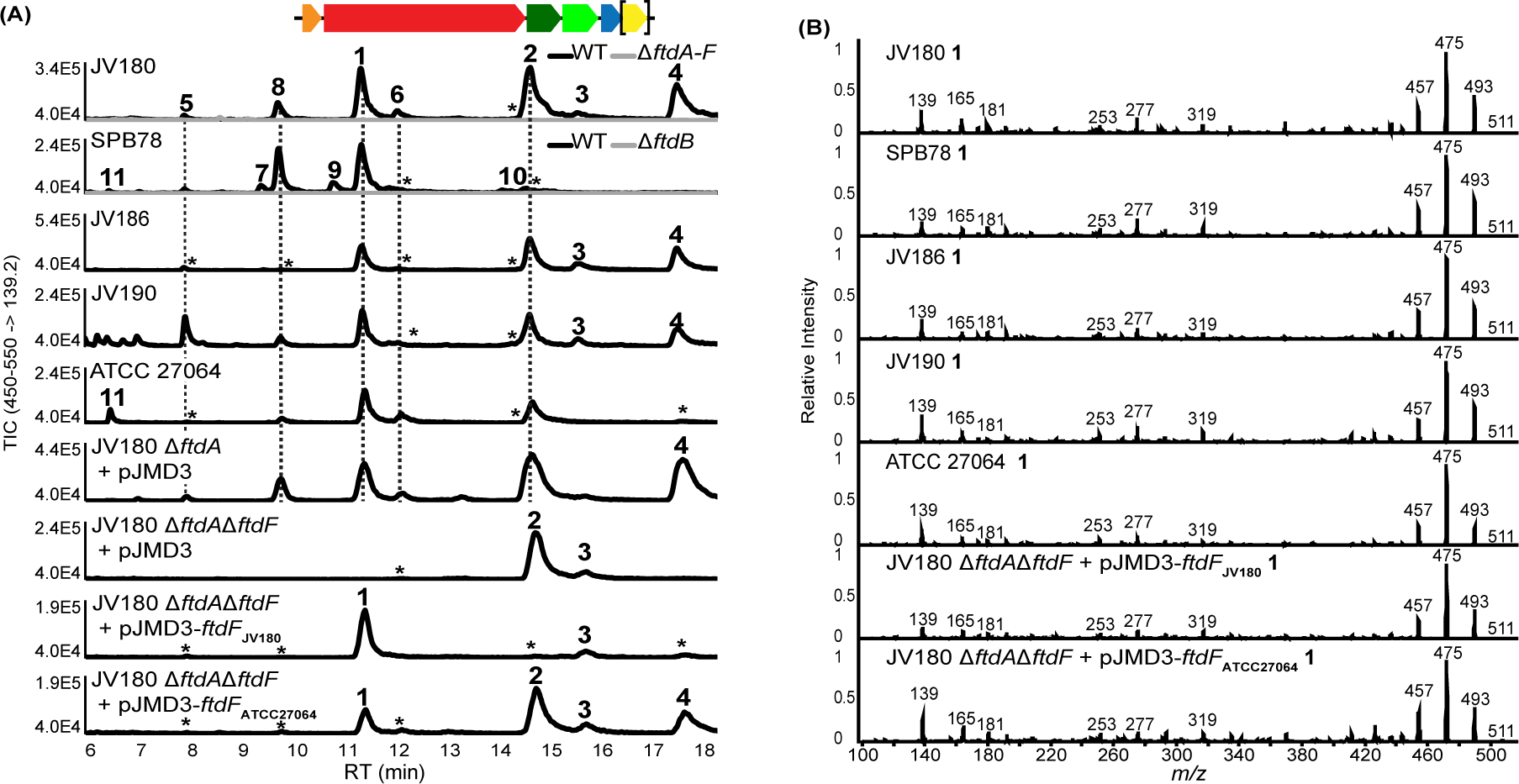
Group 1 strains produce consistent PTM congener profiles despite inconsistent *ftdF* arrangements. (A) The 25-deOH PTM products (LC-MS/MS monitoring *m/z* 139.2 product ions) of multiple Group 1 BGC strains showed largely overlapping profiles compared to PTM- mutant controls (in grey; JV168 and JV352). *ftdF*-dependent mass features were identified by comparing these against JV180 Δ*ftdA* and Δ*ftdAftdF* mutants (each with empty vector pJMD3; JV3003 and JV3006). These mutants and data were used to test for CYP450 orthology between an unclustered *S. clavuligerus ftdF* homolog (FtdF_ATCC27064_, JV2997) and ectopic native *ftdF* (FtdF*JV180*, JV3010). Asterisks indicate probable but very low abundance PTMs. (B) MS^2^ spectra of 1 from several Group 1 strains and *ftdF* mutant complements, supporting structural equivalency. The *m/z* transitions for numbered peaks are listed in Table S4.

FtdF-type CYP450s are active when their cognate genes from *S. griseus*, *S. albidoflavus*, and *Streptomyces* sp. SPB78 are heterologously expressed (22), so we expected PTM structural consequences for *S. clavuligerus*, which lacks a typical Group 1 clustered *ftdF* (clade **I** CYP450). To probe why the latter strain still produces typical Group 1 PTM mass features, we first explored *ftdF*-dependent biosynthesis in *Streptomyces* sp. JV180, a genetically amenable host. This was done by creating two mutants lacking cluster-linked oxidative genes: *Streptomyces* sp. JV180 D*ftdA* (JV1520) and double mutant D*ftdA*D*ftdF* (JV1522) (Fig 5). Mutants in *ftdA* give simplified congener diversity (2), which increases intermediate detection sensitivity by reducing congener complexity. We thus revealed compounds **1**, **4**, **5**, **8** are *ftdF*-dependent, and these were clearly present in *S. clavuligerus* growth extracts (Fig 5). This indicated the *S. clavuligerus* genome must harbor a functional *ftdF* ortholog, so BLAST was used to search the *S. clavuligerus* genome for FtdF-similar proteins along with CYP450 phylogeny to further interrogate potential hits (Fig S3). One CYP450 (69% identity to FtdF_JV180_; EFG04254.1/ FtdF_ATCC27064_) stood out as a potential ortholog, which was encoded on the 1.8MB linear plasmid pSCL4 (72).

EFG04254.1 was cloned to constitutive-expression vector pJMD3 and tested for its ability to rescue *ftdF-*dependent mass features in JV180 D*ftdA*D*ftdF* (in parallel with a FtdF_JV180_ control). After analyzing the complemented strains (JV2997 and JV3010, respectively), we found both CYP450 genes were functional in restoring production of **1** (FI-2), **4**, **5**, and **8** (Fig 5A). The major restored mass feature, **1**, was further scrutinized to confirm its equivalency to products of parent strains via LC/MS and MS/MS fragmentation (Fig 5B). JV3010 gave low production of **2** compared to the D*ftdA* and WT strains, but this may be a function of strong-constitutive expression (6). Linking EFG04254.1 to native *S. clavuligerus* PTM biosynthesis remains to be tested, but our heterologous rescue data show it catalyzes the appropriate reaction, making it a high-confidence candidate.

### Group 1 and Group 4 comparative-minimization experiments expose weaknesses in synteny-based molecule prediction

The mutational analysis of JV180 Group 1 biosynthesis revealed product congener profiles can be effectively “collapsed” to a single prominent mass-product after deleting *ftdA* and *ftdF*. We questioned what would happen if we performed a parallel analysis with other gene clusters sharing the same core BGC architecture with Group 1. Would we recover the same biosynthetic intermediate? The Group 4 BGC of *Streptomyces olivaceus* NRRL B-3009 was selected to answer this because its core BGC outwardly matches JV180, with the exception that the Group 4 strain encodes a predicted Rieske 2Fe-2S protein instead of a FtdF-type CYP450. *S. olivaceus* NRRL B-3009 PTMs were previously unanalyzed, but our PTM-targeted metabolomics detected likely products via precursor ion scans, typical UV absorbance profiles, and certain PTM-type MS product ions (Fig 6, and Fig S9C-I). As expected for strains having distinct BGC types, the putative PTM profiles of JV180 and B-3009 did not significantly overlap (Fig 6).

**Figure 6.**
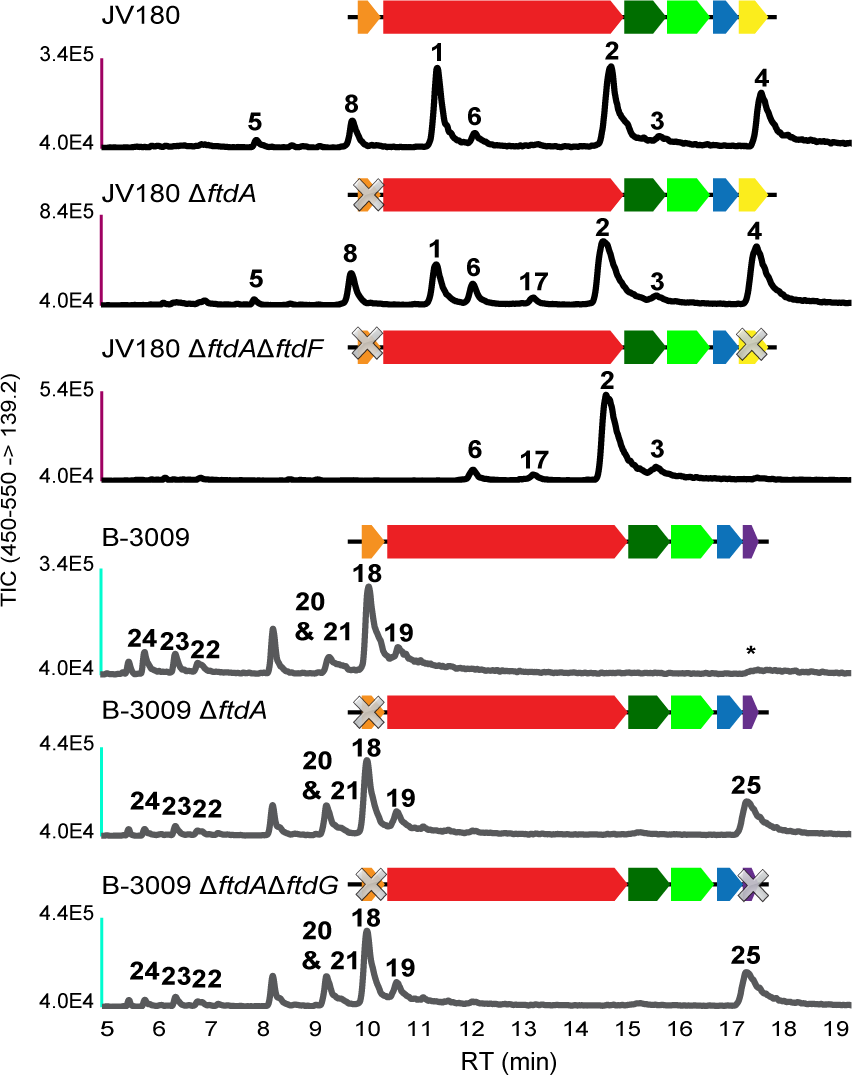
Native and tailoring-minimized Group 4 (NRRL B-3009) and Group 1 (JV180) BGCs produce group-specific product profiles. LC-MS/MS monitoring of 25-deOH PTM profiles (*m/z* 139.2 product ions) produced by *Streptomyces* sp. JV180 (JV307), JV180 Δ*ftdA* (JV1520), JV180 Δ*ftdA.ftdF* (JV1522), S. *olivaceus* NRRL B-3009 (JV1101), B-3009 Δ*ftdA* (JV1521), and B-3009 Δ*ftdA.ftdG* (JV1560) revealed unique product profiles between the tested BGC groups. Asterisks indicate PTMs were detected with certain MS/MS methods, but only in very low amounts. The *m/z* transitions for numbered peaks are listed in Table S4.

To create simplified PTM profiles in *S. olivaceus*, unmarked D*ftdA* and D*ftdA*D*ftdG* mutants were created (JV1521 and JV1560) and analyzed. Like JV180, *ftdA* mutants were expected to cease 25-OH PTM production, which was evidenced in the complete loss of *m/z* 154.2 product ion signals (Fig S9D). Surprisingly, we observed no differences in the *m/z* 139.2 precursor profiles for either mutant when compared to parent controls (Fig 6), indicating *ftdG* may not contribute to PTM biosynthesis. This was further investigated via a D*ftdG* single mutant (JV1208), and like the other mutants, this strain also lacked obvious MS/MS profile differences compared to parent controls (Fig S9C). We thus concluded *ftdG* is inessential for PTM biosynthesis under the conditions tested; any function remains unknown.

The PTM intermediates produced by the “tailoring-minimized” JV180 D*ftdA*D*ftdF* and B-3009 D*ftdA*D*ftdG* strains were then compared via LC/MS/MS, and like their wild-type parents, these mutants again failed to produce overlapping profiles (Fig 6). This suggests the existence of yet-unrecognized contributors to PTM diversity, either hidden within the genes of the PTM BGCs themselves or in host factors outside of their cognate BGCs. Supporting the former idea, the NRRL B-3009 locus is similar to that for pactamides, molecules that lack C14 hydroxyl decorations (but feature other, yet-unexplained oxidative modifications (18, 35)). Indeed, we detected a PTM signal (**31**) in NRRL B-3009 extracts which fragments consistently with a C14-deOH analog of **2** (Fig S9E-G), perhaps identical to pactamide A (*m/z* 481.3). It is thus highly plausible that NRRL B-3009 utilizes a non-hydroxylating PprB, which would partially inform why the Group 1 and 4 “tailoring minimized” strains do not accumulate the same intermediate(s). However, this cannot completely explain the differences in product profiles; the B-3009 double mutant produces several other PTM-like mass features (**18**-**30**) in addition to **31**, but these too did not match any JV180 congeners (Fig 6, S9C, S9H-J), pointing to other mechanisms of pathway divergence. These data emphasize the need to disambiguate the complex origins of these molecules, especially in terms of extra-BGC enzymology (an increasingly encountered phenomenon (73–75).

Lastly, the LC-MS/MS-based methods used here cannot resolve stereochemistry, a parameter with profound consequences on bioactivity. In light of recent PTM configuration revisions, Jin *et al* have argued for systematic re-investigation of PTM stereochemistry (24). Doing this would undoubtedly shed useful insights into structure, and eventually, refined bioinformatics models. Despite the limitations of MS, metabolomic studies on microbial chemistry are burgeoning (76–79). Because PTM BGCs are fairly common, their products are frequently encountered in these studies, but sometimes incorrectly. For example, specific PTMs have been extrapolated from MS signals whose structures are incongruent with the BGC content (i.e., butremycin from a Group 1 BGC) (80–82). Others indicate both 5/5/6 and 5/6/5 PTMs being produced by individual strains, which is unlikely without supporting genomic evidence (83–86). BGC information is thus urged as a priority dimension of these types of analyses.

## METHODS

### Strains, primers, plasmids, enzymes, chemicals, and general methods

Strains used in metabolomics analysis are listed in Table S5; primers and plasmids used in strain construction are listed in Tables S6 and S7. All primers were purchased from Integrated DNA Technologies. All restriction enzymes were purchased from New England Biolabs. Taq polymerase and T4 Ligase were purchased from New England Biolabs or Thermo Fisher Scientific. Novagen KOD Hot Start polymerase was purchased from Millipore Sigma and used for all cloning. FailSafe PCR 2X PreMix D or G (LCG, Biosearch Technologies) were used as buffer for all PCR reactions.

### Environmental actinomycete isolation and cultivation

All bacterial isolates used in this study were enriched from a diversity of soils obtained from the Tyson Research Center of Washington University in St Louis, located in Eureka, MO. Isolates *Kitasatospora* sp. strains BE20 and BE303, and *Streptomyces* sp. strains BE133, BE147, BE230, BE282, and BE308 originated from New Rankin Cave sediments. *Kitasatospora* sp. strain SP17BM10 and *Streptomyces* sp. strain SP18ES09 were isolated from grassy rhizosphere soils of New Pond field. *Streptomyces* sp. SP17KL33 (6) and SP18CS02 were isolated from dry glade environments. Enrichment media used to obtain these strains were Humic-acid Vitamin (HV), Arginine Glycerol-Salts(AGS), and Low Tryptone-Yeast (LTY), and enrichments were performed as previously described using air-dried and finely sifted soils(33). All media contained antifungals cycloheximide and nystatin, (50mg/L or 100 mg/L ea) plus either nalidixic acid (25 mg/L) or polymyxin B (10 mg/L) to discourage non-actinomycete microorganisms. Specific enrichment media for each isolate were as follows: HV for strains BE133, BE147, BE230, BE282, BE303, and BE308; AGS for strains BE20 and SP18CS02; and LTY for SP18ES09. Following isolation, the actinobacterial isolates were grown with vigorous shaking in 50mL conical tubes equipped with 5mL Trypticase Soy Broth and stored at −80°C for further use.

### Draft genome sequencing of PTM BGC-harboring environmental isolates

PTM^+^ environmental isolates above were identified using an established *ftdB*-targeted degenerate PCR screen (33). Screening amplicons were confirmed via Sanger sequencing at Genewiz using amplification primers YQ158/YQ159 (6). Parent strains of interest were draft sequenced using Illumina Genome sequencing at the Washington University in St. Louis McDonnell Genome Institute, with the TruSeq PCR-free library prep kit (Illumina) and the Sci clone next-generation sequencing (NGS) instrument (Perkin Elmer) essentially as previously reported (87).

### PTM BGC mining

AntiSMASH software versions 3, 4, and 5 were used to identify PTM BGCs among fragmentary assembled genomes from Blodgett Lab environmental isolates (88). PTM BGCs were also systematically mined from online sequence databanks including GenBank (National Center for Biotechnological Information, NCBI) and the Joint Genome Institute Integrated Microbial Genomes and Microbiomes (JGI IMG) using BLASTP and TBLASTN tools. For this, the FtdB sequence from Streptomyces sp. JV178 (54) was used as a query against all available bacterial finished, permanent, and draft genome sequences. Hits above ∼ 40% identity were selected as being potentially associated with PTM clusters, and PTM BGC identities were established through the identification of nearby encoded homologs of other known PTM BGC-associated genes. In addition, all candidate FtdB sequences were tested for expected ornithine-specific adenylation domains using the PKS/NRPS Analysis website from The Institute for Genome Sciences at U of Maryland and NRPSsp Non-Ribosomal Peptide Synthase substrate predictor (89, 90). BGC linear comparisons were visualized using Clinker (91) where necessary.

### Phylogenetic analyses of PTM BGC enzymes

For this study, the FtdB and cytochrome p450 trees were created with alignments of all available complete CDS translation products available from GenBank, plus those sequenced from environmental isolates. Specifically, these analyses included a total of 302 FtdBs and 232 CYP450s manual curated to remove exact duplicates. To ensure the quality of the FtdB tree, some FtdB having incomplete or faulty assembly were removed (including all <90% of typical FtdB length as determined by the average the lengths of 6 published FtdB’s, plus 6 FtdB’s sampling diverse genera found in this study (Table S1). In contrast, the FtdA, PhyDH, and ADH trees were created with enzyme sequences extracted from representative strains drawn from each BGC group (see Fig 2 for BGC group definitions). This was done for figure clarity and to high sequence redundancy. Sequences were aligned using Qiagen CLC Main Workbench v20. Maximum Likelihood trees were created using IQ-TREE2 with the model finder plus setting to determine the best-fitting model and 1000 ultra-fast bootstrapping replicates to assess branch supports. Ultrafast bootstrap threshold was set at 95%, and branches over 99% were bolded for each tree. Trees were visualized in CLC Main Workbench. The best-fit models selected for each tree are listed: FtdA: mtInv+F+I+G4, FtdB: JTT+F+R10, FtdB amplicon: JTT+F+R10, Ppr: LG+F+R7, ADH: LG+F+R5, Cyp450: LG+F+R5.

Specific outgroups to root each phylogeny were: for PhyDH (IkaB/FtdC), carotene dehydratase family enzyme from *Rhodobacter capsulatus* (CrtI) (CAA77540.1) from (2); for cytochrome p450s, *Bacillus megaterium* BM3 (WP_095379236.1) from (6); for FtdB an iterative Type 1 PKS enzyme from *Saccharopolyspora erythraea* (WP_009944722.1) was used. To identify suitable outgroups to root ADH and FtdA trees, examples of each PTM associated protein were analyzed using BlastP (NCBI) to identify associated protein superfamilies. This revealed candidate non-PTM associated members for each superfamily. For the PTM ADH tree, an oxidoreductase from *Bacillus subtilis* subsp. *subtilis* str. 168 (CAB12574.1) was selected, and for FtdA, the root is a sterol desaturase from *Agrobacterium fabrum* str. C58 (Q7D1P6).

### Taxonomic classification PTM BGC-harboring strains

PTM BGC-harboring genomes were downloaded from NCBI, as well as select partial housekeeping sequences (*atpD, recA, rpoB,* and *trpB*) of type strains used in previous taxonomic studies (92). Automlsa2 was used with model finder plus and 1000 ultrafast bootstrap replicates to mine and align sequences, and then build a phylogenetic tree (Best-fit model: GTR+F+R8:atpD+recA,GTR+F+R5:rpoB,TVM+F+R6:trpB) (93). CLC Main Workbench v20.0 was used to visualize alignments and trees. Published phylogenies using MLSA, whole genome analysis, and ANI scores were taken into consideration when analyzing taxonomic classification (92, 94–101).

### PTM production and targeted metabolomics

Strains were revived from −80°C cryostocks onto ISP2 agar and incubated 2-3 d at 28°C. Agar plugs with adhering mycelia were used to innoculate single colonies to 3 ml TSB in 24-well plates. Cultures were shaken for 2 days at 28°C (250 rpm) with a sterile 4mm glass bead to disrupt clumping. 200 µl of each culture was spread to ISP2, ISP4, ISPS, SFM, or YMS agar plates and incubated at 28°C for 6 days. The agar was diced, immersed in ethyl acetate overnight, and solvent was evaporated under vacuum. The dry extracts were resuspended in 500 µl LC-MS grade methanol and filtered before HPLC analysis.

Chromatography was performed using a Phenomenex Luna C18 column (75 x 3 mm, 3 µm pore size) installed on an Agilent 1260 Infinity HPLC. For each run, 10 µl of sample was injected and the chromatography conditions were as follows: T = 0, 5% B; T = 1, 5% B; T = 4, 35% B; T = 22, 55% B; T = 26, 100% B; T = 30, 100%; A: water + 0.1% formic acid, B: acetonitrile + 0.1% formic acid; and 0.8 ml/min. An HPLC diode array detector (DAD) monitored PTM production at 320 nm, along with in-line Agilent 6420 Triple-Quad mass spectrometer for MS/MS analysis with ESI in positive mode (ESI+). For PTM-targeted metabolomics, precursor ion scan mode was used to detect molecules *m/z* 450 – 550 that fragmented (collision energy = 30 V, fragmentation energy = 70 V) into product ions *m/z* 139.2 and 154.2 as previously (38). Product ion scan mode was then used to detect precursor ions that were identified in the precursor scans, *m/z*: 493.2, 495.4, 497.4, 509.2, 511.2, 513.2, 525.2, 527.3, 529.0. MS data were analyzed offline using Agilent MassHunter Qualitative Analysis software, where PTMs were identified by correlating baseline-corrected UV profiles with PTM-like fragmentation patterns, established in (2, 6, 38).

### Genetics: Marker-less gene deletion and rescue experiments

Streptomycin counterselections were used to construct unmarked mutants to test gene function essentially as described (6, 38). Briefly, protein S12 (StrR) point mutants of *S. olivaceus* NRRL B-3009 were isolated by plating cells to ISP2 + Str^10^, and *rpsL* was amplified from resistant colonies and sequenced using primers YQ4 and YQ5 (Table S7). The *rpsL* mutants used as parents for downstream mutation analyses were chosen based on their having had no visible difference in PTM production or growth characteristics from WT. All gene deletions were generated using double homologous recombination as previously described (6, 38). Deletion plasmids were generated by first cloning PCR-amplified homologous upstream and downstream regions into linearized pUC19 via NEB Hi-Fi Assembly and verified by Sanger sequencing. Homology cassettes were subcloned to pJVD52.1. Resulting plasmids were introduced into *Streptomyces* via intergeneric conjugation using JV36, where apramycin-resistant (Apr^R^), Str^S^ exconjugants were selected. These were grown in TSB at 37°C without antibiotics, and double recombinants were identified via PCR after plating to ISP2 + streptomycin. Mutant rescue and heterologous expression were carried out using expression-shuttle vector pJMD3. Inserts were confirmed by Sanger sequencing using primers P*ermE*\*-fw and PXS6 or via whole plasmid sequencing by Plasmidsaurus using Oxford Nanopore Technologies. Expression constructs were conjugated into recipients using JV36 by selecting for Apr^R^ colonies, essentially as previously described (2). Plasmid integration at FC31 *attB* was confirmed using multiplex PCR as done (6). See Table S6 for specific plasmid details.

### FtdB Clade-based primer design and function tests

A total of 11 degenerate primer sets (see Table S3) were designed based *ftdB* nucleotide sequences drawn from distinct phylogenetic groups of the FtdB ML-tree (Fig S1). Degenerate primers were designed to specifically amplify across the junction of the PKS and NRPS modules of the enzymes using Primers4Clades(102). Primers were designed using the default *get primers* run mode, and with corrected CODEHOP (codeh_corr) formulations. A set of positive control strains (Table S3) were selected to validate the functionality of the new degenerate primer pairs. *Streptomyces* sp. SP18CS02 has a newly-described cluster architecture, and it was also used to test new primers because the strain saw inconsistent amplification with YQ158/YQ159 (6). Universal 16s rDNA primers 8F/1492R (103) were used to control for template quality. All new degenerate *ftdB* screening primers were used with the following “touchdown” protocol: 2 min at 95 °C, followed by 7 cycles consisting of 20 s at 95 °C, 15 s at 70 °C, and 1 min at 70 °C, decreasing annealing temperature by 2 °C after each cycle. This was followed by 23 cycles consisting of 20 s at 95 °C, 15 s at 54 °C, and 1 min at 68 °C, ending with 5 min at 72 °C. All primer sets yielded expected amplicon sizes based on clade-specific positive-control *ftdB* gene sequences (see Table S3).

### Nucleotide accession numbers

The Whole Genome Shotgun Sequencing data have deposited at DDBJ/ENA/GenBank under accessions: JAQYWT000000000 for *Streptomyces sp.* SP18BB07, JAQYWU000000000 for *Streptomyces sp.* BE133, JAQYWV000000000 for *Streptomyces sp.* BE147, JAQYWW000000000 for *Streptomyces sp.* BE20, JAQYWX000000000 for *Streptomyces sp.* BE230, JAQYWY000000000 for *Streptomyces sp.* BE282, JAQYWZ000000000 for *Streptomyces sp.* BE303, JAQYXA000000000 for *Streptomyces sp.* BE308, JAQYXB000000000 for *Streptomyces sp.* SP17BM10, JAQYXC000000000 for *Streptomyces sp.* SP18CS02, JAQYXD000000000 for *Streptomyces sp.* SP18ES09, JAQYXE000000000 for *Streptomyces sp.* JV176, JAQYXF000000000 for *Streptomyces sp.* JV181, JAQYXG000000000 for *Streptomyces sp.* JV184, JAQYXH000000000 for *Streptomyces sp.* JV185, JAQYXI000000000 for *Streptomyces sp.* JV186, JAQYXJ000000000 for *Streptomyces sp.* JV190, and JAQYXK000000000 for *Streptomyces sp.* SP17KL33.

## SUPPLEMENTAL MATERIALS

Fig S1 to S9 and Table S1 to S7.

## ACKNOWLEDGMENTS

We thank John D’Alessandro, Brandon Burger, Eric Song, Bilal Makhdoom, and Carrie Stump for assisting strain isolations, Priya Mullick for cloning assistance, and Catrina Fronick for draft genome assembly. For strains or DNA to validate degenerate PCR toolsets, we thank: Paul Jensen (University of California-San Diego; *Salinispora arenicola* CNS-205;), Liangcheng Du (University of Nebraska-Lincoln; *Lysobacter enzymogenes* C3), and Avena Ross (Queen’s University-Ontario; *Pseudoalteramonas luteoviolacea* 2ta16). This work was supported by NSF-CAREER 1846005 to J.A.V.B. Undergraduate researcher support is also gratefully acknowledged to: E.Z (Amgen Scholars Award), A. D and M.T (Washington University Biology Summer Undergraduate Research Fellowships). M.T also received a Lynda A. Ceremsak & F. George Davitt Undergraduate Biochemistry Fellowship and a WUSTL-Howard Hughes Medical Institute Summer Undergraduate Research Fellowship. J.A.V.B received genome-sequencing support through a McDonnell Genome Institute 2018 Symposium Pilot Project funded by the Office of the Dean of the School of Medicine, Washington University Institute of Clinical and Translational Sciences (ICTS), and Illumina, Inc. (San Diego, CA).

