## Supplementary Information for "Critical analysis of polycyclic tetramate macrolactam biosynthetic cluster phylogeny and functional diversity"

[d]USDA-ARS, 1100 Robert E. Lee Blvd. New Orleans, LA 70124

**Contents**

**Figure S1.** Maximum-likelihood phylogenetic tree generated from 302 full-length FtdB sequences.

**Figure S2**. Maximum-likelihood phylogenetic tree made from a concatenated alignment of four partial housekeeping gene sequences (*atpD*, *recA*, *rpoB*, and *trpB*) estimates the relatedness of strains.

**Figure S3**. Maximum-likelihood phylogenetic tree generated from 232 CYP450 sequences.

**Figure S4**. Maximum-likelihood phylogenetic tree generated from representative FtdA sequences.

**Figure S5.** Maximum-likelihood phylogenetic tree generated from representative ADH sequences.

**Figure S6.** Genetic locus plasticity contributes to PTM BGC diversity in Group 4.

**Figure S7.** PCR amplicons of representative strains generated with newly designed primers help capture more *ftdB* diversity.

**Figure S8**. Comparison of FtdB amplicon sequences vs full-length FtdB sequences.

**Figure S9**. Group 1 strains produce consistent PTMs congener profiles, which are different than PTM congener profile of Group 4 strain B-3009.

**Table S1.** FtdB lengths.

**Table S2.** Strains used in FtdB phylogeny and their assigned BGC groups.

**Table S3.** Clade-based *ftdB* degenerate primers.

**Table S4.** *m/z* transitions for each PTM identified in metabolomics analysis.

**Table S5.** Strains used for cloning, sequencing, and metabolomics.

**Table S6.** Plasmids used in strain construction.

**Table S7.** Primers used in strain construction.

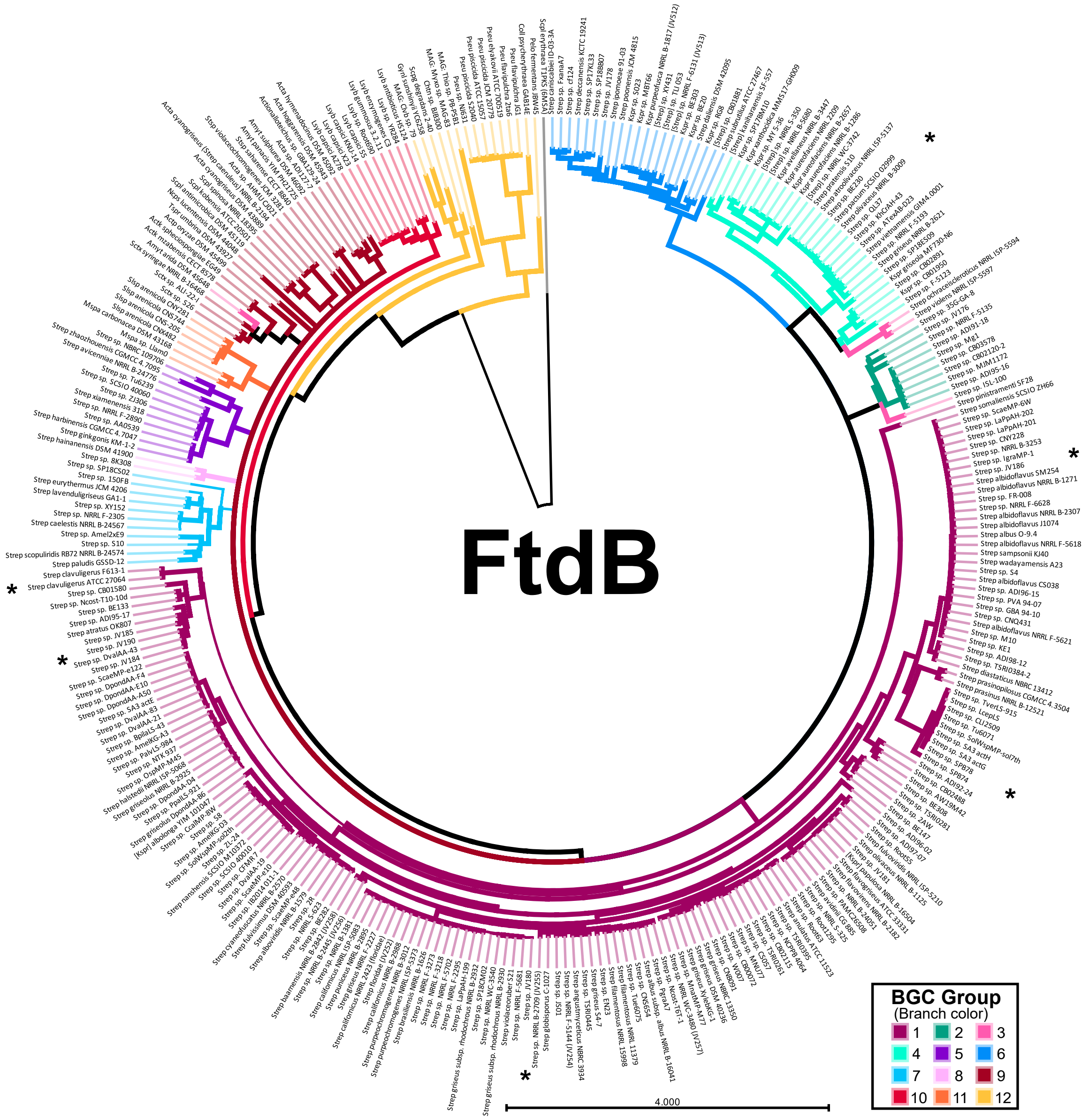


**Fig S1** Maximum-likelihood phylogenetic tree generated from 302 full-length FtdB sequences. Branches are colored according to BGC groups. UF-Bootstrap threshold is 95%, branches above 99% are bolded. Asterisks indicate strains used in metabolomics.

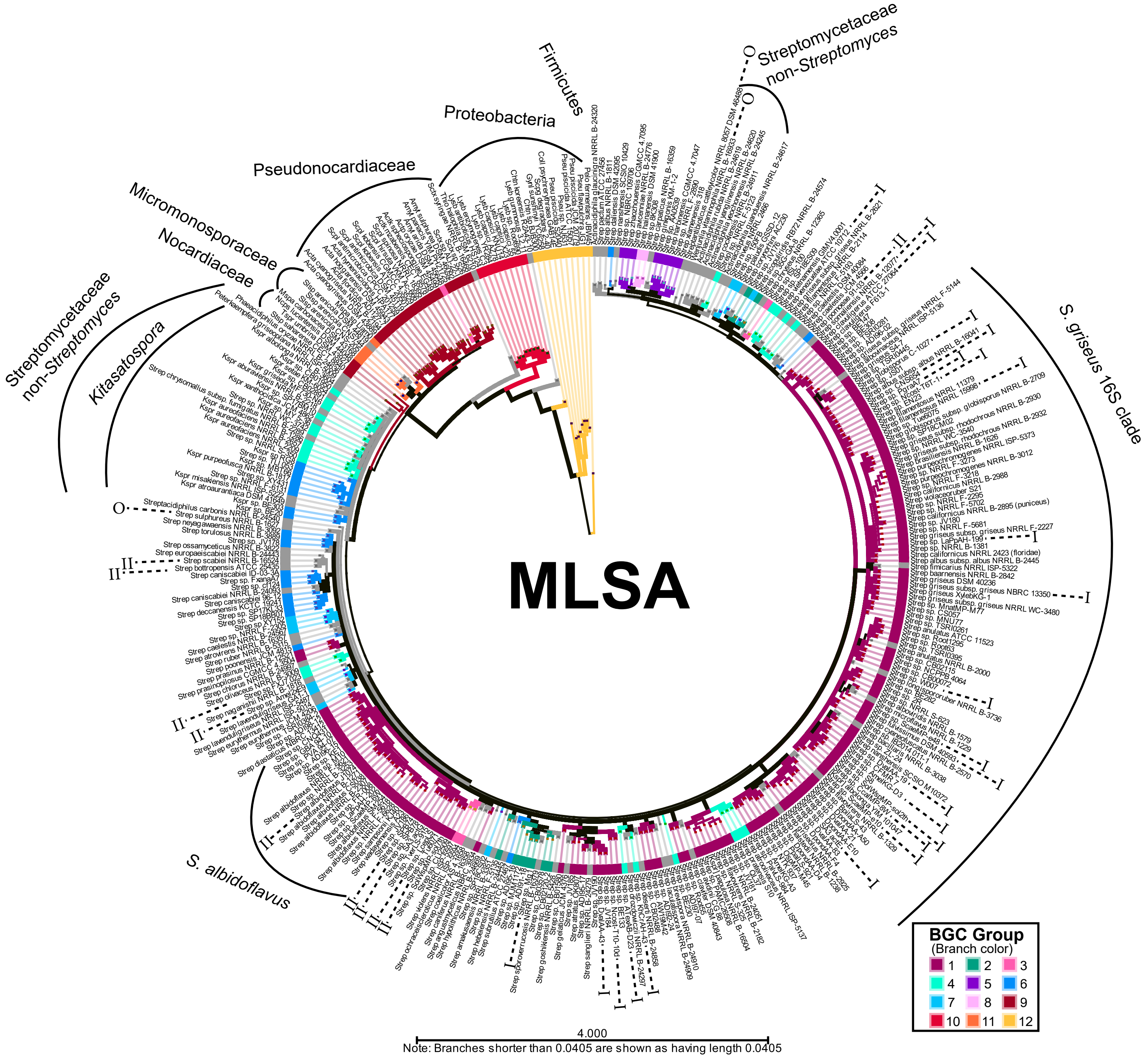


**Fig S2** Maximum-likelihood phylogenetic tree made from a concatenated alignment of four partial housekeeping gene sequences (*atpD*, *recA*, *rpoB*, and *trpB*) estimates the relatedness of strains. Branches are colored according to BGC groups; grey branches are reference strains. UF-Bootstrap threshold is 95%, branches above 99% are bolded. I, II, and O (other) designate *Streptomyces* lineages as defined by McDonald and Currie (2017).

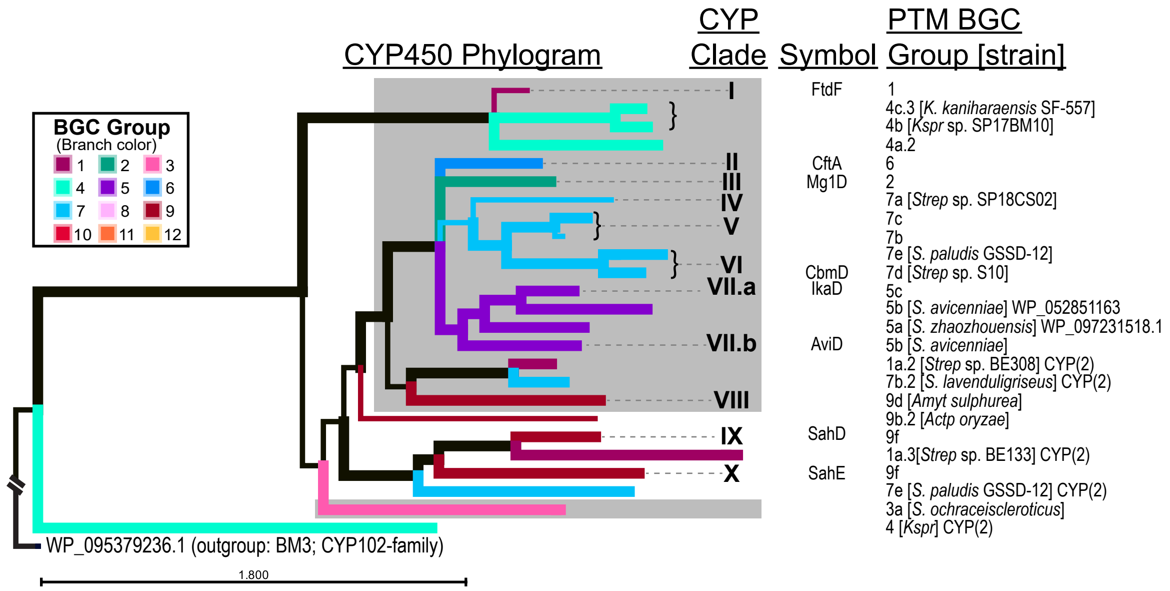


**Fig S3** Maximum-likelihood phylogenetic tree generated from 232 CYP450 sequences. Branches are colored according to BGC groups. Shading indicates CYP107-family enzymes, which are known to be polyphyletic (2). UF-Bootstrap threshold is 95%, branches above 99% are bolded.

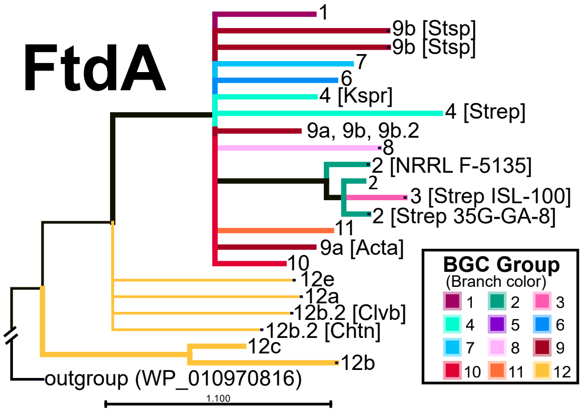


**Fig S4** Maximum-likelihood phylogenetic tree generated from representative FtdA sequences. Branches are colored according to PTM groups. Branches are collapsed where all leaves are part of the same BGC group. UF-Bootstrap threshold is 95%, branches above 99% are bolded.

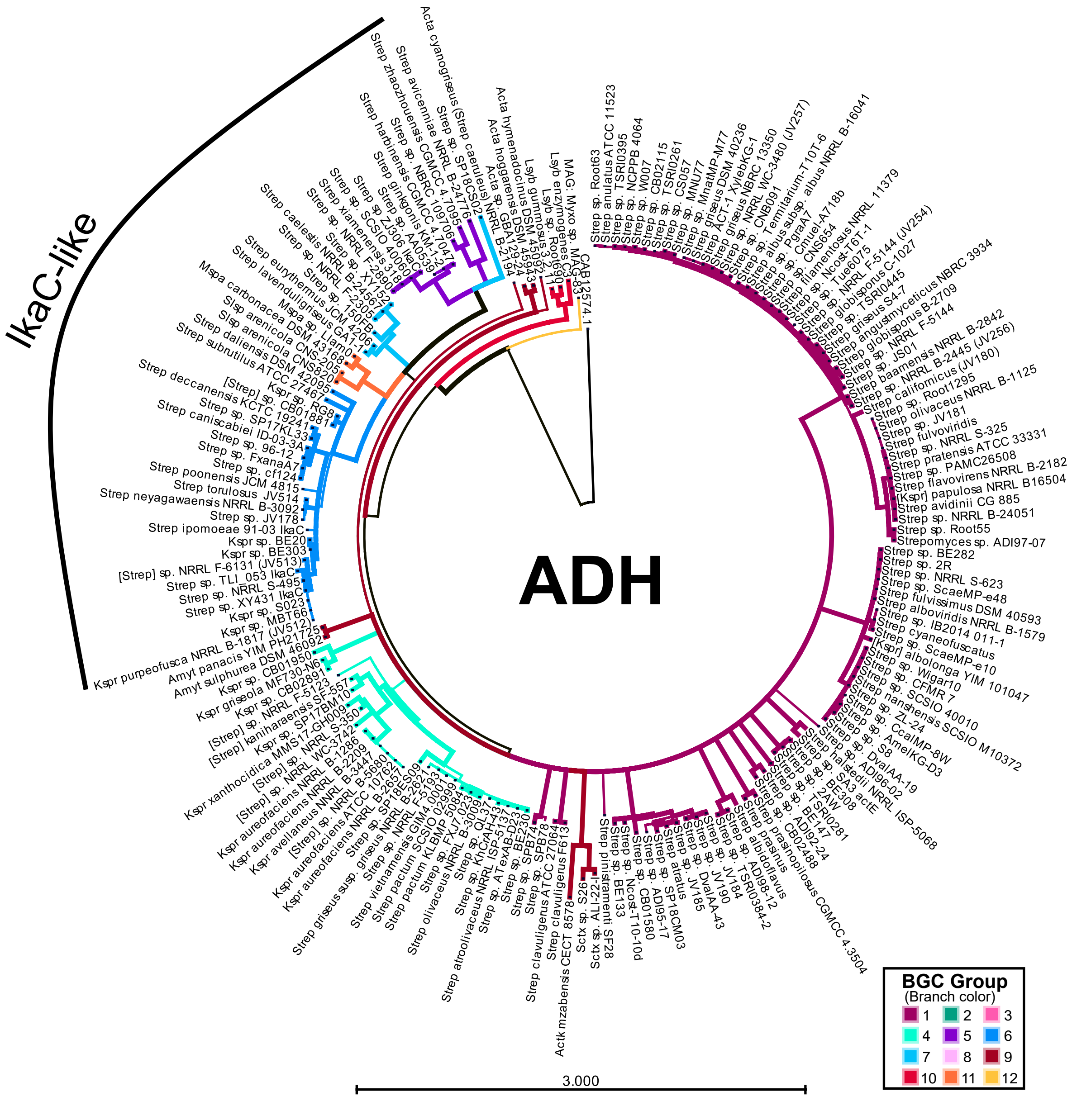


**Fig S5** Maximum-likelihood phylogenetic tree generated from representative C-ring forming ADH sequences. Branches are colored according to PTM groups. UF-Bootstrap threshold is 95%, branches above 99% are bolded.

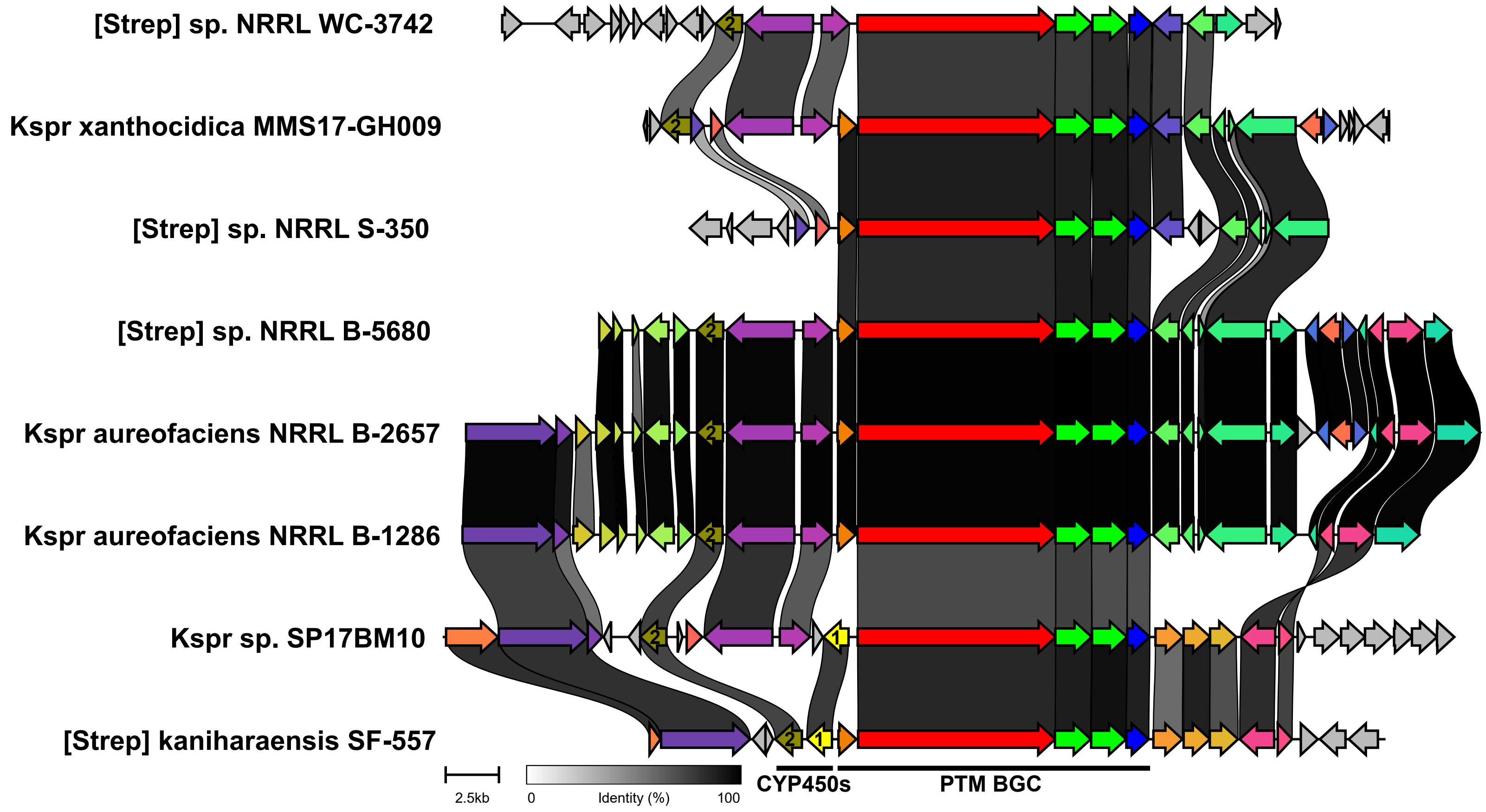


**Fig S6** Genetic locus plasticity contributes to PTM BGC diversity in Group 4. Genes with significant similarity are colored the same.

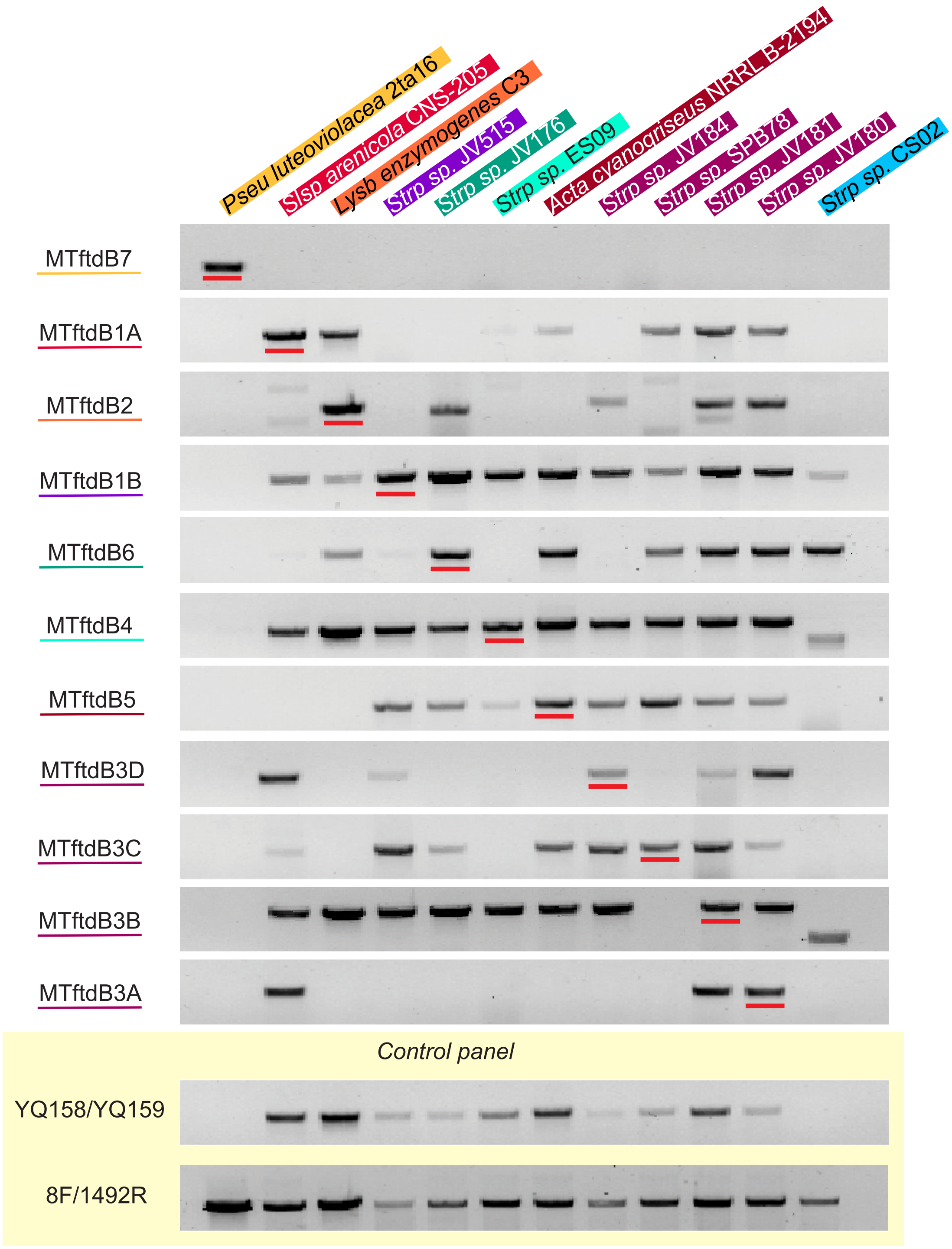


**Fig S7** PCR amplicons of representative strains generated with newly designed primers help capture more *ftdB* diversity. Amplicons from the representative strain of each primer set are underlined in red. Amplicons from previously published *ftdB* primers and universal 16S primers are shown in control panel.

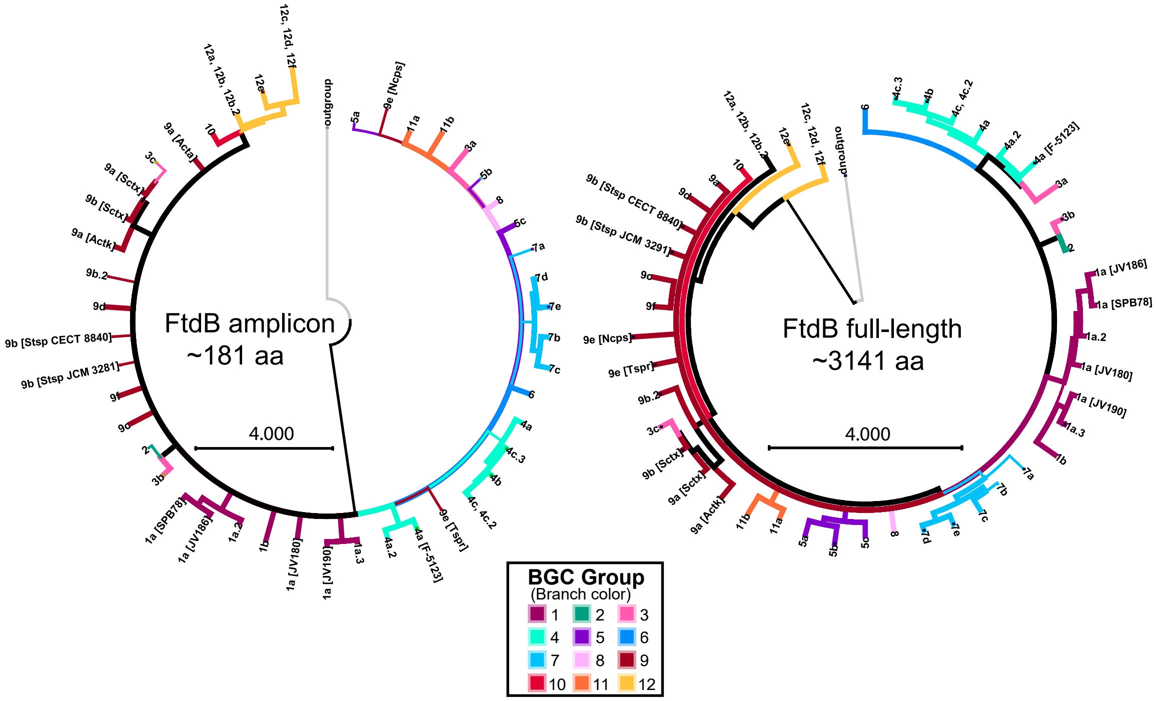


**Fig S8** Phylogenetic comparison of FtdB amplicons vs full-length FtdB sequences reveals that novelty can be estimated from short internal fragments of *ftdB*. Maximum-likelihood phylogenetic trees generated from 302 (A) FtdB amplicon sequences (~181 aa) amplified using primers YQ158 and YQ159 and (B) FtdB full-length sequences (~3141 aa). UF-Bootstrap threshold is 95%; Branches above 99% are bolded.

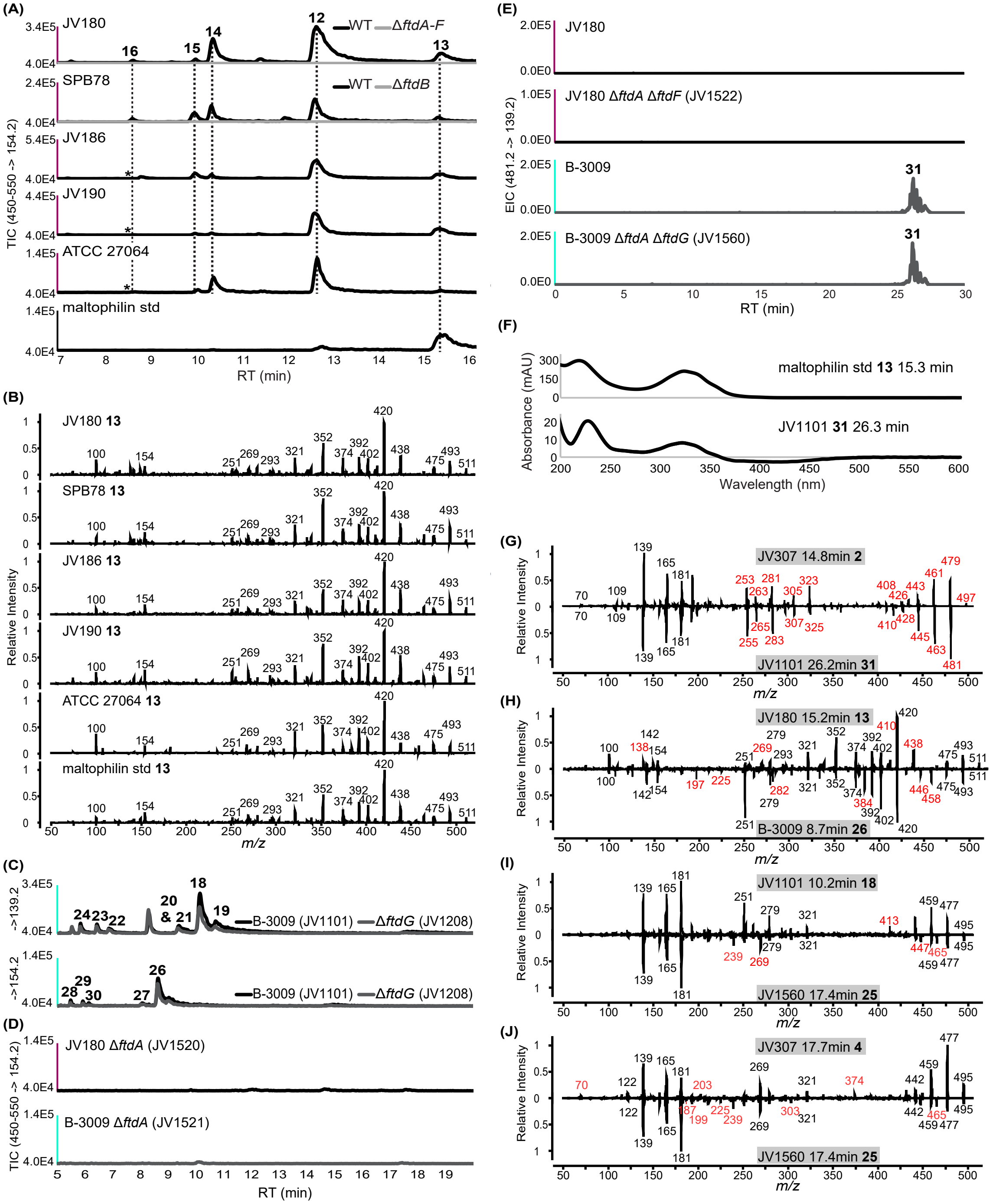

**FIG S9** Group 1 strains produce consistent PTM congener profiles, which are different than the PTM congener profile of Group 4 strain B-3009. (A) LC-MS/MS monitoring of 25-OH PTMs (*m/z* 154.2 product ions) from Group 1 strains and null PTM mutants. (B) $MS^2$ spectra of PTM **13** from Group 1 strains and a maltophilin standard. (C) LC-MS/MS monitoring of PTMs (*m/z* 139.2, 154.2 product ions) shows that *ftdG* is not essential for PTM biosynthesis. (D) Δ*ftdA* mutants of JV180 or B-3009 do not produce 25-OH PTMs, as expected. (E) B-3009 produces a PTM **31** with the same *m/z* as pactamide A. (F) PTM **31** exhibits a PTM-like UV spectrum. (G) $MS^2$ mirror plot of **2** vs **31**. (H) $MS^2$ mirror plot of **13** vs **26**. (I) $MS^2$ mirror plot of **18** vs **25**. (J) $MS^2$ mirror plot of **4** vs **25**. Asterisks indicate PTMs were detected with certain MS/MS methods, but only in very low amounts. The *m/z* transitions for numbered peaks are listed in Table S4.

**Table S1:** Calculations of the average amino acid length of FtdB using 6 canonical FtdB's and 6 FtdB's from this study that were from diverse genera to create a well-rounded data set. The average amino acid length of all 12 enzyme sequences was calculated, and then the "cutoff" for new FtdB sequences for the FtdB tree was determined as being 90% of the average length.

| Strain name | Amino acid length (AA) | Average amino acid length (AA) | Cutoff for AA length (AA) |
|---|---|---|---|
| Lysobacter enzymogenes C3 (3) | 3123 | 3141.3 | 2827.2 |
| Streptomyces sp. SPB78 (4) | 3138 | | |
| Streptomyces xiamenensis 318 (5) | 3124 | | |
| Streptomyces sp. JV178 (6) | 3284 | | |
| Streptomyces sp. KL33 (6) | 3150 | | |
| Streptomyces albus J1074 (7) | 3137 | | |
| Colwellia psycherythraea GAB14E | 3138 | | |
| Actinoalloteichus cyanogriseus DSM 43889 | 3124 | | |
| Salinspora arenicola CNS-205 | 3101 | | |
| Actinokineospora spheciospongiae EG49 | 3048 | | |
| Nocardiopsis lucentensis DSM 44048 | 3216 | | |
| Pseudoalteromonas sp. NJ631 | 3113 | | |

**Table S2.** Strains used for FtdB phylogeny and their assigned PTM BGC Group.

| Abbreviated Name | Group |
|---|---|
| Strp caniscabiei ID-03-3A | 6a |
| Strp sp. FxanaA7 | 6a |
| Strp sp. cf124 | 6a |
| Strp deccanensis KCTC 19241 | 6a |
| Strp sp. SP17KL33 | 6a |
| Strp sp. SP18BB07 | 6a |
| Strp sp. JV178 | 6b |
| Strp ipomoeae 91-03 | 6b |
| Strp poonensis JCM 4815 | 6a |
| Kspr sp. S023 | 6b |
| Kspr sp. MBT66 | 6b |
| Kspr purpeofusca NRRL B-1817 (JV512) | 6b |
| [Strp] sp. XY431 | 6b |
| [Strp] sp. TLI 053 | 6b |
| [Strp] sp. NRRL F-6131 (JV513) | 6b |
| Kspr sp. BE303 | 6b |
| Kspr sp. BE20 | 6b |
| Strp daliensis DSM 42095 | 6c |
| Kspr sp. RG8 | 6b |
| [Strp] sp. CB01881 | 6b |
| Strp subrutilus ATCC 27467 | 6b |
| [Strp] kaniharaensis SF-557 | 4c.3 |
| Kspr sp. SP17BM10 | 4b |
| Kspr xanthocidica MMS17-GH009 | 4c.2 |
| Kspr sp. MY 5-36 | 4c.2 |
| [Strp] sp. NRRL S-350 | 4c |
| [Strp] sp. NRRL B-5680 | 4c.2 |
| Kspr avellaneus NRRL B-3447 | 4c.2 |
| Kspr aureofaciens NRRL 2209 | 4c.2 |
| Kspr aureofaciens NRRL B-2657 | 4c.2 |
| Kspr aureofaciens NRRL B-1286 | 4c.2 |
| [Strp] sp. NRRL WC-3742 | 4b.2 |
| Strp pratensis S10 | 4a |
| Strp atroolivaceus NRRL ISP-5137 | 4a |
| Strp sp. BE230 | 4a |
| Strp pactum SCSIO 02999 | 4a |
| Strp olivaceus NRRL B-3009 | 4a |
| Strp sp. QL37 | 4a |
| Strp sp. KhCrAH-43 | 4a |
| Strp sp. ATexAB-D23 | 4a |
| Strp vietnamensis GIM4.0001 | 4a |
| Strp sp. NRRL F-5193 | 4a |
| Strp griseus NRRL B-2621 | 4a |
| Strp sp. SP18ES09 | 4a |
| Kspr griseola MF730-N6 | 4a.2 |
| Strp sp. CB02891 | 4a.2 |
| Kspr sp. CB01950 | 4a.2 |
| Strp sp. F-5123 | 4a |
| Strp ochraceiscleroticus NRRL ISP-5594 | 3a |
| Strp violens NRRL ISP-5597 | 3a |
| Strp sp. 35G-GA-8 | 2 |
| Strp sp. JV176 | 2 |
| Strp sp. NRRL F-5135 | 2 |
| Strp sp. ADI91-18 | 2 |
| Strp sp. Mg1 | 2 |
| Strp sp. CB03578 | 2 |
| Strp sp. CB02120-2 | 2 |
| Strp sp. MJM1172 | 2 |
| Strp sp. ADI95-16 | 2 |
| Strp sp. ISL-100 | 3b |
| Strp pinistramenti SF28 | 1a |
| Strp somaliensis SCSIO ZH66 | 1a |
| Strp sp. ScaeMP-6W | 1a |
| Strp sp. LaPpAH-202 | 1a |
| Strp sp. LaPpAH-201 | 1a |
| Strp sp. CNY228 | 1a |
| Strp sp. NRRL B-3253 | 1a |
| Strp sp. IgraMP-1 | 1a |
| Strp sp. JV186 | 1a |
| Strp albidoflavus SM254 | 1a |
| Strp albidoflavus NRRL B-1271 | 1a |
| Strp sp. FR-008 | 1a |
| Strp sp. NRRL F-6628 | 1a |
| Strp albidoflavus NRRL B-2307 | 1a |
| Strp albidoflavus J1074 | 1a |
| Strp albus O-9.4 | 1a |
| Strp albidoflavus NRRL F-5618 | 1a |
| Strp sampsonii KJ40 | 1a |
| Strp wadayamensis A23 | 1a |
| Strp sp. S4 | 1a |
| Strp albidoflavus CS038 | 1a |

| | |
|---|---|
| Strp sp. ADI96-15 | 1a |
| Strp sp. PVA 94-07 | 1a |
| Strp sp. GBA 94-10 | 1a |
| Strp sp. CNQ431 | 1a |
| Strp albidoflavus NRRL F-5621 | 1a |
| Strp sp. M10 | 1a |
| Strp sp. KE1 | 1a |
| Strp sp. ADI98-12 | 1a |
| Strp sp. TSRI0384-2 | 1a |
| Strp diastaticus NBRC 13412 | 1a |
| Strp prasinopilosus CGMCC 4.3504 | 1a |
| Strp prasinus NRRL B-12521 | 1a |
| Strp sp. TverLS-915 | 1a |
| Strp sp. LcepLS | 1a |
| Strp sp. CLI2509 | 1a |
| Strp sp. Tu6071 | 1a |
| Strp sp. SolWspMP-sol7th | 1a |
| Strp sp. SA3 actH | 1a |
| Strp sp. SA3 actG | 1a |
| Strp sp. SPB78 | 1a |
| Strp sp. SPB74 | 1a |
| Strp sp. ADI92-24 | 1a |
| Strp sp. CB02488 | 1a |
| Strp sp. AW19M42 | 1a |
| Strp sp. BE308 | 1a.2 |
| Strp sp. TSRI0281 | 1a.2 |
| Strp sp. 2AW | 1a.2 |
| Strp sp. BE147 | 1a.2 |
| Strp sp. ADI96-02 | 1a |
| Strp sp. ADI97-07 | 1a |
| Strp sp. Root55 | 1a |
| Strp fulvoviridis NRRL ISP-5210 | 1a |
| Strp olivaceus NRRL B-1125 | 1a |
| Strp sp. JV181 | 1a |
| [Kspr] papulosa NRRL B-16504 | 1a |
| Strp flavogriseus ATCC 33331 | 1a |
| Strp flavovirens NRRL B-2182 | 1a |
| Strp sp. NRRL B-24051 | 1a |
| Strp sp. PAMC26508 | 1a |
| Strp avidinii CG 885 | 1a |
| Strp sp. NRRL S-325 | 1a |
| Strp sp. Root1295 | 1a |
| Strp sp. Root63 | 1a |

| | |
|---|---|
| Strp anulatus ATCC 11523 | 1a |
| Strp sp. TSRI0395 | 1a |
| Strp sp. NCPPB 4064 | 1a |
| Strp sp. CB02115 | 1a |
| Strp sp. TSRI0261 | 1a |
| Strp sp. CS057 | 1a |
| Strp sp. MNU77 | 1a |
| Strp sp. CB00072 | 1a |
| Strp sp. W007 | 1a |
| Strp sp. CNB091 | 1a |
| Strp griseus NBRC 13350 | 1a |
| Strp griseus DSM 40236 | 1a |
| Strp griseus XylebKG-1 | 1a |
| Strp sp. MnatMP-M77 | 1a |
| Strp sp. NRRL WC-3480 (JV257) | 1a |
| Strp sp. Ncost-T6T-1 | 1a |
| Strp sp. PgraA7 | 1a |
| Strp albus subsp. albus NRRL B-16041 | 1a |
| Strp sp. CNS654 | 1a |
| Strp sp. Tue6075 | 1a |
| Strp filamentosus NRRL 11379 | 1a |
| Strp filamentosus NRRL 15998 | 1a |
| Strp sp. EN23 | 1a |
| Strp griseus S4-7 | 1a |
| Strp sp. TSRI0445 | 1a |
| Strp angustmyceticus NBRC 3934 | 1a |
| Strp sp. NRRL F-5144 (JV254) | 1a |
| Strp sp. JS01 | 1a |
| Strp globisporus C-1027 | 1a |
| Strp sp. NRRL B-2709 (JV255) | 1a |
| Strp sp. JV180 | 1a |
| Strp sp. NRRL F-5681 | 1a |
| Strp violaceoruber S21 | 1a |
| Strp griseus subsp. rhodochrous NRRL B-2930 | 1a |
| Strp sp. NRRL WC-3540 | 1a |
| Strp sp. SP18CM02 | 1a |
| Strp griseus subsp. rhodochrous NRRL B-2932 | 1a |
| Strp sp. LaPpAH-199 | 1a |
| Strp sp. NRRL F-2295 | 1a |
| Strp sp. NRRL F-5702 | 1a |
| Strp sp. NRRL F-3218 | 1a |

| | |
|---|---|
| Strp sp. NRRL F-3273 | 1a |
| Strp brasiliensis NRRL B-1626 | 1a |
| Strp purpeochromogenes NRRL ISP-5373 | 1a |
| Strp purpeochromogenes NRRL B-3012 | 1a |
| Strp californicus NRRL B-2988 | 1a |
| Strp floridae (JV252) | 1a |
| Strp californicus NRRL 2423 (floridae) | 1a |
| Strp griseus NRRL F-2227 | 1a |
| Strp puniceus NRRL B-2895 | 1a |
| Strp californicus NRRL ISP-5083 | 1a |
| Strp sp. NRRL B-1381 | 1a |
| Strp sp. NRRL B-2445 (JV256) | 1a |
| Strp baarnensis NRRL B-2842 (JV258) | 1a |
| Strp sp. BE282 | 1a |
| Strp sp. NRRL S-623 | 1a |
| Strp sp. 2R | 1a |
| Strp alboviridis NRRL B-1579 | 1a |
| Strp sp. ScaeMP-e48 | 1a |
| Strp fulvissimus DSM 40593 | 1a |
| Strp cyaneofuscatus NRRL B-2570 | 1a |
| Strp sp. IB2014 011-1 | 1a |
| Strp sp. ScaeMP-e10 | 1a |
| Strp sp. DvalAA-19 | 1a |
| Strp sp. CFMR 7 | 1a |
| Strp sp. SCSIO 40010 | 1a |
| Strp nanshensis SCSIO M10372 | 1a |
| Strp sp. ZL-24 | 1a |
| Strp sp. SolWspMP-sol2th | 1a |
| Strp sp. AmelKG-D3 | 1a |
| Strp sp. S8 | 1a |
| Strp sp. CcalMP-8W | 1a |
| [Kspr] albolonga YIM 101047 | 1a |
| Strp griseolus DpondAA-B6 | 1a |
| Strp sp. PpalLS-921 | 1a |
| Strp sp. DpondAA-D4 | 1a |
| Strp griseolus NRRL B-2925 | 1a |
| Strp halstedii NRRL ISP-5068 | 1a |
| Strp sp. OspMP-M45 | 1a |
| Strp sp. NTK 937 | 1a |
| Strp sp. PalvLS-984 | 1a |
| Strp sp. AmelKG-A3 | 1a |

| | |
|---|---|
| Strp sp. BpilaLS-43 | 1a |
| Strp sp. DvalAA-21 | 1a |
| Strp sp. DvalAA-83 | 1a |
| Strp sp. SA3 actE | 1a |
| Strp sp. DpondAA-A50 | 1a |
| Strp sp. DpondAA-E10 | 1a |
| Strp sp. DpondAA-F4 | 1a |
| Strp sp. ScaeMP-e122 | 1a |
| Strp sp. JV184 | 1a |
| Strp sp. DvalAA-43 | 1a |
| Strp sp. JV190 | 1a |
| Strp sp. JV185 | 1a |
| Strp atratus OK807 | 1a |
| Strp sp. ADI95-17 | 1a |
| Strp sp. BE133 | 1a.3 |
| Strp sp. Ncost-T10-10d | 1a.3 |
| Strp sp. CB01580 | 1a.3 |
| Strp clavuligerus ATCC 27064 | 1a |
| Strp clavuligerus F613-1 | 1a |
| Strp paludis GSSD-12 | 7e |
| Strp scopuliridis RB72 NRRL B-24574 | 7d |
| Strp sp. S10 | 7d |
| Strp sp. Amel2xE9 | 7d |
| Strp caelestis NRRL B-24567 | 7c |
| Strp sp. NRRL F-2305 | 7c |
| Strp sp. XY152 | 7c |
| Strp lavenduligriseus GA1-1 | 7b |
| Strp eurythermus JCM 4206 | 7b |
| Strp sp. 150FB | 7b |
| Strp sp. SP18CS02 | 7a |
| Strp sp. 8K308 | 8 |
| Strp hainanensis DSM 41900 | 8 |
| Strp ginkgonis KM-1-2 | 5c |
| Strp harbinensis CGMCC 4.7047 | 5c |
| Strp sp. AA0539 | 5c |
| Strp sp. NRRL F-2890 | 5c |
| Strp xiamenensis 318 | 5c |
| Strp sp. ZJ306 | 5c |
| Strp sp. SCSIO 40060 | 5c |
| Strp sp. Tu6239 | 5c |
| Strp avicenniae NRRL B-24776 | 5b |
| Strp zhaozhouensis CGMCC 4.7095 | 5a |

| | |
|---|---|
| Strp sp. NBRC 109706 | 5a |
| Mspa sp. Llam0 | 11a |
| Mspa carbonacea DSM 43168 | 11a |
| Slsp arenicola CNX482 | 11b |
| Slsp arenicola CNS-205 | 11b |
| Slsp arenicola CNS744 | 11b |
| Slsp arenicola CNY281 | 11b |
| Sctx sp. S26 | 9a |
| Sctx sp. ALI-22-I | 9a |
| Sctx syringae NRRL B-16468 | 9b |
| Amyt arida DSM 45648 | 3c |
| Actk mzabensis CECT 8578 | 9a |
| Actk spheciospongiae EG49 | 9a |
| Actp oryzae DSM 45499 | 9b.2 |
| Tspr umbrina DSM 43927 | 9e |
| Ncps lucentensis DSM 44048 | 9e |
| Scpl antimicrobica DSM 45119 | 9f |
| Scpl kobensis ATCC 20501 | 9f |
| Scpl spinosa NRRL 18395 | 9f |
| Acta cyanogriseus DSM 43889 | 9c |
| Acta cyanogriseus (Strp caeruleus) NRRL B-2194 | 9c |
| Acta sp. AHMU CJ021 | 9c |
| Stsp violaceochromogenes JCM 3281 | 9b |
| Stsp saharense CECT 8840 | 9b |
| Amyt panacis YIM PH21725 | 9d |
| Amyt sulphurea DSM 46092 | 9d |
| Acta sp. ADI127-7 | 9a |
| Acta sp. GBA129-24 | 9a |
| Acta hoggarensis DSM 45943 | 9a |
| Acta hymenadocinus DSM 45092 | 9a |
| Lsyb capsici AZ78 | 10a |
| Lsyb capsici X23 | 10a |
| Lsyb capsici KNU-14 | 10a |
| Lsyb capsici 55 | 10a |
| Lsyb sp. Root690 | 10a |
| Lsyb gummosus 3.2.11 | 10a |
| Lsyb enzymogenes C3 | 10a |
| Lsyb sp. YR284 | 10a |
| Lsyb antibioticus HS124 | 10b |
| MAG: Clvb sp. 79 | 12b.2 |
| Gynl sunshinyii YC6258 | 12b |
| Scpg degradans 2-40 | 12a |

| | |
|---|---|
| Chtn sp. BJB300 | 12b.2 |
| MAG: Myxo sp. MAG-83 | 12e |
| MAG: Thio sp. PB-PSB1 | 12f |
| Pseu sp. NJ631 | 12c |
| Pseu piscicida S2040 | 12c |
| Pseu piscicida ATCC 15057 | 12c |
| Pseu piscicida JCM 20779 | 12c |
| Pseu elyakovii ATCC 700519 | 12c |
| Pseu flavipulchra 2ta6 | 12c |
| Pseu flavipulchra JG1 | 12c |
| Coll psycherythraea GAB14E | 12c |
| Pelo fermentans JBW45 | 12d |

Abbreviations:
Strp = Streptomyces
Kspr = Kitasatospora
Mspa = Micromonospora
Slsp = Salinispora
Sctx = Saccharothrix
Amyt = Amycolatopsis
Actk = Actinokineospora
Actp = Actinophytocola
Tspr = Thermomonospora
Ncps = Nocardiopsis
Scpl = Saccharopolyspora
Acta = Actinoalloteichus
Stsp = Streptosporangium
Lysb = Lysobacter
Clvb = Cellvibrio
Gynl = Gynuella
Scpg = Saccharophagus
Chtn = Chitinimonas
Myxo = Myxococcales (order)
Thio = Thiohalocapsa
Pseu = Pseudoalteromonas
Coll = Colwellia
Pelo = Pelosinus
MAG = Metagenome Assembled Genome

**Table S3.** Clade-based *ftdB* degenerate primers.

| **Primer set** | **Represented group(s)** | **Template strains (GenBank accession)** | **Average amplicon size** | **Positive control** |
|---|---|---|---|---|
| MTftdB1A-F/ MTftdB1A-R | 11a, 11b | GCA_000375065.1, GCA_900091535.1 | 650 bp | *Salinispora arenicola* CNS-205 |
| MTftdB1B-F/ MTftdB1B-R | 6a, 6b, 5a, 5b, 5c | GCA_000710405.2, GCA_900230195.1, GCA_000974485.1, GCA_900105395.1, GCA_000719915.1, JAQYXK000000000, CP009922.3, CP026498.1, MF893273.1, KF954512.1 | 750 bp | *Streptomyces sp.* JV515 |
| MTftdB2-F/ MTftdB2-R | 10a, 10b | GCA_900107375.1, GCA_000336385.3, GCA_000604005.2, GCA_002355295.1, GCA_001427785.1, LBMI00000000, CP040656.1, CP011130.1, CP023465.1 | 500 bp | *Lysobacter enzymogenes* C3 |
| MTftdB3A-F/ MTftdB3A-R | 1a | GCA_014748265.1, GCA_001279425.1, GCA_000156455.1, GCA_000719585.1, GCA_900090135.1, GCA_900187925.1, GCA_000932225.2, CP031425.1 | 500 bp | *Streptomyces sp*. JV180 |
| MTftdB3B-F/ MTftdB3B-R | 1a | GCA_002705725.1, GCA_900091865.1, GCA_002242735.1, GCA_000382745.1, GCA_003259275.1, GCA_003259575.1, JJOB00000000, CP020563.1, CP002993.1 | 750 bp | *Streptomyces sp.* JV181 |
| MTftdB3C-F/ MTftdB3C-R | 1a | GCA_900091485.1, GCA_002802945.1, GCA_000721455.1, GCA_000800535.1, CP014485.1, CP016824.1, CP110818.1 | 700 bp | *Streptomyces sp.* SP78 |
| MTftdB3D-F/ MTftdB3D-R | 1a, 1a.3 | GCA_900091955.1, JAQYXG000000000, JAQYXJ000000000, JAQYXH000000000, JAQYWU000000000 | 500 bp | *Streptomyces sp.* JV184 |
| MTftdB4-F /MTftdB4-R | 4a, 4a.2 | GCA_000721515.1, GCA_000836635.1, GCA_000721495.1, GCA_000373645.1, GCA_000373445.1, GCA_000373505.1, JAQYWX000000000, JAQYXD000000000 | 650bp | *Streptomyces sp.* SP18ES09 |
| MTftdB5-F/ MTftdB5-R | 4b, 4b.2, 4c, 4c.2, 9a, 9c, 9d, | GCA_002564045.1, GCA_000429185.2, GCA_000720635.1, GCA_000719705.1, GCA_000719415.1, JAQYXB000000000, CP025990.1, CP014859.1, CP022521.1 | 700 bp | *Actinoalloteichus cyanogriseus* NRRL B-2194 |
| MTftdB6-F/MTftdB6-R | 2, 7b, 7c, 7d | GCA_000718375.1, GCA_000818195.1, GCA_000718095.1, GCA_000719775.1, JAQYXE000000000, CP011664.1 | 700 bp | *Streptomyces sp.* JV176 |
| MTftdB7-F/MTftdB7-R | 12a, 12b, 12c, 12d | GCA_000495575.1, GCA_000967555.1, GCA_000382005.1, GCA_000764185.1, GCA_000259115.1, JX173670.1, CP011924.1, CP000282.1, CP010978.1, BK010667.1 | 800 bp | *Pseudoalteromonas luteoviolacea* 2ta16 |

<u>MTftdB1A</u>
F: CGCTGGACCACTTCgtvctsttygc
R: AGCTGCTTGAGGaaccasarygc
Amplicon size: 768 bp

MTftdB1B
F: CGCTGGACCACTTCgtvctsttcgc
R: ACCACAGGGCCTTCtggttstgsgt
Amplicon size: 765 bp

MTftdB2
F: CCGCCGGCCAGrccaactaygc
R: CGGCGCCGccgatgttgta
Amplicon size: 765 bp

MTftdB3A
F: CCTCCTCGCCatggarctrcg
R: GGGGGTCGTGGGcsagrtcgtasg
Amplicon size: 498 bp

MTftdB3B
F: CGCTGGACCACTTCgtsctsttcgc
R: CCGGGTTCAGGTGCttsaggaacca
Amplicon size: 777 bp

MTftdB3C
F: AGACCAACTACGCCgcsggsaacgc
R: CACAGGGCCTTCtggttctssgt
Amplicon size: 702 bp

MTftdB3D
F: GGAATGATCGAAGAACTAGGACTArtcgrycacta
R: CCTGAGGAACCATAGAGCTTTCtggttctssgt
Amplicon size: 562 bp

MTftdB4
F: CGTGGGCCACCGGcatgatcgagg
R: GGCGCCGCCGatgttgtasgc
Amplicon size: 660 bp

MTftdB5
F: AGACCAACTACGCCgcsggsaacgc
R: ACCACAGGGCCTTCtggttstgbgt
Amplicon size: 702 bp

MTftdB6
F: CGCCTTCCTAGACGCTCTAgcccaccaccg
R: TTGTTTGAGGAACCATAGAGCCttctggttctg
Amplicon size: 723 bp

MTftdB7B
F: CGGCAGACATATAGCCCAAT
R: ATGGAGATGGTGCGAATGAT
Amplicon size: 412 bp

**Table S4.** *m/z* transitions for each PTM identified in this study (Fig 5-6, Fig S7).

| PTM | *m/z* transition | notes |
|---|---|---|
| 1 | 511->139 | FI-2 |
| 2 | 497->139 | |
| 3 | 495->139 | 280/380 UV max |
| 4 | 495->139 | |
| 5 | 513->139 | |
| 6 | 495->139 | |
| 7 | 509->139 | |
| 8 | 509->139 | FI-1 |
| 9 | 511->139 | |
| 10 | 493->139 | |
| 11 | 511->139 | |
| 12 | 513->154 | |
| 13 | 511->154 | maltophilin-like |
| 14 | 511->154 | |
| 15 | 527->154 | |
| 16 | 525->154 | frontalamide A |
| 17 | 497->139 | |
| 18 | 495->139 | |
| 19 | 495->139 | |
| 20 | 495->139 | |
| 21 | 509->139 | |
| 22 | 513->139 | |
| 23 | 513->139 | |
| 24 | 513->139 | |
| 25 | 495->139 | |
| 26 | 511->154 | |
| 27 | 511->154 | |
| 28 | 529->154 | |
| 29 | 529->154 | |
| 30 | 529->154 | |
| 31 | 481->139 | |

**Table S5**. Strains used for cloning, sequencing, and metabolomics.

| Strain | Genotype/ characteristics | Source |
|---|---|---|
| DH5α λ*pir* | *E. coli* cloning host, F– φ80lacZΔM15 Δ(lacZYA-argF)U169 recA1 endA1 hsdR17 (rK–, mK+) phoA supE44 λ– thi-1 gyrA96 relA1 | NEB |
| JV36 | *E. coli* conjugal donor strain, dam-3 dcm-6 metB1 galK2 galT27 lacY1 tsx-78 supE44 thi-1 mel-1 tonA31 ΔhsdRMS-mrr::FRT(rK-mK-) attHK::pJK202 (ΔoriR6K-aadA::FRT bla::pir) (Tra+, AmpS) | (4) |
| *Streptomyces* sp. JV180 | Wild type | (4) |
| *Streptomyces* sp. SPB78 | Wild type | (4) |
| *Streptomyces* sp. JV186 | Wild type | (4) |
| *Streptomyces* sp. JV190 | Wild type | (4) |
| *S. clavuligerus* ATCC 27064 | Wild type | ATCC |
| *S. olivaceus* NRRL B-3009 | Wild type | NRRL |
| *Streptomyces* sp. JV176 | Wild type | (4) |
| *Streptomyces* sp. JV181 | Wild type | (4) |
| *Streptomyces* sp. JV184 | Wild type | (4) |
| *Streptomyces* sp. JV185 | Wild type | (4) |
| *Streptomyces* sp. SP17KL33 | Wild type | (6) |
| *Streptomyces* sp. SP18BB07 | Wild type | This study |
| *Streptomyces* sp. SP17BM10 | Wild type | This study |
| *Streptomyces* sp. SP18CS02 | Wild type | This study |
| *Streptomyces* sp. SP18ES09 | Wild type | This study |
| *Streptomyces* sp. BE133 | Wild type | This study |
| *Streptomyces* sp. BE147 | Wild type | This study |
| *Streptomyces* sp. BE20 | Wild type | This study |
| *Streptomyces* sp. BE230 | Wild type | This study |
| *Streptomyces* sp. BE282 | Wild type | This study |
| *Streptomyces* sp. BE303 | Wild type | This study |
| *Streptomyces* sp. BE308 | Wild type | This study |
| JV307 | JV180 *rpsL*_K43R | (9) |
| JV352 | JV307 Δ*ftdA-F* | (9) |

| | | |
|---|---|---|
| JV141 | SPB78 *rpsL*_K43R | (4) |
| JV168 | JV141 Δ*ftdB* | (4) |
| JV1101 | B-3009 *rpsL*_K43R | This study |
| JV1208 | JV1101 *ΔftdG* | This study |
| JV1520 | JV307 Δ*ftdA* | This study |
| JV1521 | JV1101 Δ*ftdA* | This study |
| JV1522 | JV307 Δ*ftdA ΔftdF* | This study |
| JV1560 | JV1101 Δ*ftdA ΔftdG* | This study |
| JV2997 | JV1522 attBΦC31::pJMD3-ftdF$_{ATCC27064}$ | This study |
| JV3003 | JV1520 attBΦC31::pJMD3 | This study |
| JV3006 | JV1522 attBΦC31::pJMD3 | This study |
| JV3010 | JV1522 attBΦC31::pJMD3-ftdF$_{JV180}$ | This study |

**Table S6.** Plasmids used in strain construction.

| Plasmid | Description | Source |
|---|---|---|
| pUC19 | *bla ori*$^{pUC}$ cloning vector; used for subcloning | NEB |
| pETDuet-1 | T7 promoter expression vector; used for subcloning | Novagen |
| pJVD52.1 | Apr$^{R}$ , *oriT*, *rep*$^{pSG5(ts)}$, *rep*$^{pUC}$, *rpsL*$_{S.\ coelicolor}$ shuttle vector | (8) |
| pJMD3 | *aac(3)IV oriT* attP-*int*ΦC31 *rep*pUC self-integrating vector containing PermE*pHM11a promoter | (6, 10) |
| pUC19-deltarieske | gibson assembly of PCR products of pUC19 (primers YQ268-pUC19-us and YQ269-pUC19-ds), JV1101 (primers ED18 and ED19; ED20 and ED21) | This study |
| pJVD52.1-deltarieske | ligation of pJVD52.1 and pUC19-deltarieske, both digested with XbaI and HindIII | This study |
| pUC19-deltaftdAJV180 (pSDS005) | gibson assembly of pUC19 digested with EcoRI and XbaI and PCR products of JV180 (primers SS44 and SS45; SS46 and SS47) | This study |
| pUC19-deltaftdAJV820 (pSDS006) | gibson assembly of pUC19 digested with BamHI and XbaI and PCR products of JV1101 (primers SS52 and SS53; SS54 and SS55) | This study |
| pJVD52.1-deltaftdAJV180 (pSDS007) | ligation of pJVD52.1 and SDS005, both digested with XbaI and EcoRI | This study |
| pJVD52.1-deltaftdAJV820 (pSDS008) | ligation of pJVD52.1 and SDS006, both digested with XbaI and BamHI | This study |
| pUC19-deltaftdF | gibson assembly of PCR products of pUC19 (primers YQ268-pUC19-us and YQ269-pUC19-ds), JV1101 (primers YQ367 and YQ368; YQ84 and YQ369) | This study |
| pJVD52.1-deltaftdF | ligation of pJVD52.1 and pUC19-deltaftdF, both digested with XbaI and HindIII | This study |
| pETDuet-1-ftdF_ATCC 27064 | Ligation of PCR product of *S. clavuligerus* ATCC 27064 (primers PM07_P450_*S.clavuligerus*_F and PM08_P450_*S.clavuligerus*_R) and pETDuet, both digested with NdeI and XbaI | This study |
| pJMD3-ftdF_ATCC 27064 | Ligation of pETDuet-1-ftdF_ATCC27064 and pJMD3, both digested with NdeI and XbaI | This study |
| pJMD3-ftdF_JV180 | ligation of pJMD3 digested with Nde1 and PCR product of ftdF (YQ401 and YQ402) | This study |

**Table S7.** Primers used in strain construction.

| Primers | Sequence (5'->3') | Description |
|---|---|---|
| PermE*-fw | **GGCGCGCC**AGCCCGACCCGAGCACGC | sequencing pJMD3 insert |
| PXS6 | GGCCGATTCATTAATGCAGC | |
| R2NJ | TCGTGGCCGTCCAGCYCNCCNGCDAT | check for integration at phiC31 *attB* site |
| F3NJ | GACCCGTTCATCATGATGGAYCARATGGG | |
| Int 7NJ | CTCTTCGTTCGTCTGGAAGG | |
| Int 8NJ | CTTGTCTTCGTTGGCGCTAC | |
| YQ268-pUC19-us | TCTAGAGGATCCCCGGGTAC | linearize pUC19 |
| YQ269-pUC19-ds | AAGCTTGGCGTAATCATGGTC | |
| YQ4-Sg_rpsL_f | CACGAACGGCACACAGAAAC | sequencing *rpsL* |
| YQ5-Sg_rpsL_r | GATGATGACCGGGCGCTTC | |
| SS44-dFtdA_us_Fwd_JV180 | GTTGTAAAACGACGGCCAGTGAATTCGCTGACCACCACCTCGTCGCCCG | amplify upstream region of *ftdA* in JV180 |
| SS45-dFtda_us_rev_JV180 | CGGTGCGGGTCCCATCGGCGGATGGTTCGCGCTC | |
| SS46-dFtdA_ds_fwd_JV180 | TCCGCCGATGGGACCCGCACCGGGCACCACC | amplify downstream region of *ftdA* in JV180 |
| SS47-dFtdA_ds_rev_JV180 | CATGCCTGCAGGTCGACTCTAGACGTGCGCCTCCATGTACTGGAGGTC | |
| SS48-JV180_ftdA_US_ckF | GAAGTCGGCCATGCGCACGTTC | amplify or sequence *ftdA* deletion region in JV180 |
| SS49-JV180_ftdA_US_ckR | CCGGTTGTTCACGTCGGCCATG | |
| SS50-JV180_ftdA_DS_ckF | GACCTGCGGTGACGCGATGC | |
| SS51-JV180_ftdA_DS_ckR | CCAGGGCGTTGGCCTCGATCG | |
| SS52-dFtda_us_fwd_JV820 | TCGAGCTCGGTACCCGGGGATCCATCTACAAGAACCGGGTGACG | amplify upstream region of *ftdA* in B-3009 |
| SS53-dFtdA_us_rev_JV820 | GGCGCACGGGGGACCCCGTATCGGGTCGATC | |
| SS54-dFtdA_ds_fwd_JV820 | CGATACGGGGTCCCCCGTGCGCCGCCCGAC | amplify downstream region of *ftdA* in B-3009 |
| SS55-dFtdA_ds_rev_JV820 | CTTGCATGCCTGCAGGTCGACTCTAGAGTTGGTGCGTCCGTCCTGGTTCAC | |
| SS56-JV820_ftdA_US_ckF | CCTGTTCCTGGTCGGCGACACC | amplify or sequence *ftdA* deletion region in B-3009 |
| SS57-JV820_ftdA_US_ckR | GGTACAGCGGGCATCGCGGTAC | |
| SS58-JV820_ftdA_DS_ckF | CCGCTGCTACTCTGCGAACTCC | |
| SS59-JV820_ftdA_DS_ckR | CAGACCCGCTCGATGAGCGTGG | |
| YQ367-180DftdF-us-f | GGCCAGTGAATTCGAGCTCGGTACCCGGGGATCCTCTAGACTCGATCATGGAGGTGGTGC | amplify upstream of *ftdF* in JV180 |
| YQ368-180DftdF-us-r | TCAACCGGCTGCACGACGGGACCGTGCACGACTGGTGAGCGCTCAGAGTTCGACGAG | |
| YQ84-180PTM_ds_f | TCACCAGTCGTGCACGGTCCC | |

| | | |
|---|---|---|
| YQ369-180DftdF-ds-r | TCACACAGGAAACAGCTATGACCATGATTACGCCAAGCTTGTCGCTGTACCGGGGCGCG | amplify downstream of *ftdF* in JV180 |
| ED18 | GGCCAGTGAATTCGAGCTCGGTACCCGGGGATCC**TCTAGA**CTCGATCATGGAGGTGCTGG | amplify upstream region of *ftdG* in B-3009, includes puc19 Xba**I** enzyme site |
| ED19 | AGCGGCCGGGCTACGTCACCCTGCTCTCCGCTTCCCCCGCCGTGAGCGCCCGGGTGTGGC | amplify upstream region of *ftdG* in B-3009, includes 40 nucleotides of downstream gene |
| ED20 | GCGGGGGAAGCGGAGAGCAG | amplify downstream region of *ftdG* in B-3009 |
| ED21 | TCACACAGGAAACAGCTATGACCATGATTACGCC**AAGCTT**GAACTGGTCGAACTGGCCG | amplify downstream region of *ftdG* in B-3009, includes puc19 Hind**III** enzyme site |
| ED22 | GGCTCGCGGACTTCTATCTG | amplify or sequence *ftdG* deletion region in B-3009 |
| ED23 | TCATGACCGACACCGTCTTC | |
| ED24 | GCTTCACCTCACTCTCCCTG | |
| ED25 | GAATGAGACTCGCCGCTGTG | |
| PM07_P450_*S.clavuligerus*_F | ATATGCTCCATATGGCGCCCGGAGGGCTGCCGTCC | amplify *ftdF* in *S. clavuligerus* ATCC 27064 |
| PM08_P450_*S.clavuligerus*_R | ATATGCTCTCTAGACCTTCACGGGGGCGGGTTAATCG | |
| YQ401-180-ftdF-f | actagtgaCATATGACCACCGTCGACCCCACG | amplify *ftdF* in JV180 |
| YQ402-180-ftdF-r | actagtgaTCTAGACTACCAGGTGACTCCGAGGG | |
